# Longitudinal monitoring of individual infection progression in *Drosophila melanogaster*

**DOI:** 10.1101/2021.08.17.456698

**Authors:** Bryan A. Ramirez-Corona, Anna C. Love, Srikiran Chandrasekaran, Jennifer A. Prescher, Zeba Wunderlich

## Abstract

The innate immune system is critical for host survival of infection. Infection models in organisms like *Drosophila melanogaster* are key for understanding evolution and dynamics of innate immunity. However, current toolsets for fly infection studies are limited in their throughput and their ability to generate longitudinal measurements of infection progression in single animals. Here we report a novel bioluminescent imaging strategy enabling non-invasive characterization of pathogen load over time. We demonstrate that photon flux from autobioluminescent reporter bacteria can be used to monitor relative pathogen loads in individual animals. *Escherichia coli* expressing the *ilux* operon were imaged in whole, living flies at relevant concentrations for immune study. Because animal sacrifice was not necessary to estimate pathogen load, stochastic responses to infection were characterized in individuals over time for the first time. The high temporal resolution of bioluminescence imaging also enabled visualization of the fine dynamics of microbial clearance on the hours time-scale. Overall, this non-invasive imaging strategy provides a simple and scalable platform to observe changes in pathogen load *in vivo* over time.

## Introduction

Bacteria are widespread and can cause severe disease in animals, including humans (Hunter et al., 2010). Hosts fight infection using their immune system. Infection progression and outcome is ultimately determined by a combination of genetics, environment, and stochastic events (Carruthers et al., 2020; Duneau et al., 2017). Determining the relative contribution of each of these factors to individual prognosis will enable the identification of genetic markers and early predictors of infection outcome. Such discoveries will contribute to targeted treatment of bacterial infections.

Determination of genetic and stochastic contributions to infection outcome requires a host organism amenable to genetic manipulation and high-throughput experimentation. *Drosophila melanogaster* fulfills both of these criteria. There are thousands of inbred and sequenced *D. melanogaster* lines, and flies are tractable for high-throughput experimentation in 96-well plates, unlike common mammalian model organisms (Lack et al., 2016). Additionally, *Drosophila* possess an innate immune system composed of signaling pathways that are highly conserved in mammals (Lemaitre & Hoffmann, 2007). Flies use both the Toll and IMD signaling pathways in their immune response. The Toll pathway was initially discovered in flies and is analogous to the Toll-like receptor signaling pathway found in mammals (Lemaitre & Hoffmann, 2007). In both flies and mammals, the pathway depends on molecular recognition through pattern recognition receptors (PRRs) to initiate downstream immune response (Moy & Cherry, 2013). The IMD pathway in flies is orthologous to the TNF receptor family signaling cascade in mammals (Buchon, Silverman, & Cherry, 2014; Lemaitre & Hoffmann, 2007). Beyond their use as a platform for discovering conserved immune genes, studying immunity in insects like *Drosophila* provides insight into how insect vectored diseases spread, and how they may be contained (Schneider, 2000).

Variation in infection outcome between *D. melanogaster* lines is in part determined by the genetic backgrounds of the host (Duneau et al., 2017; Hotson and Schneider, 2015; Lazzaro et al., 2006; Sackton et al., 2010). Using genetically distinct *D. melanogaster* lines, previous work has identified how different loci affect either the ability of an animal to reduce bacterial load or to induce gene expression upon infection, for example (Frochaux et al., 2020; Lazzaro et al., 2006). A fly’s ability to endure infection -- tolerance -- is less studied both in its mechanism of action and its variation between lines. Tolerance is difficult to measure in high throughput, as it requires measuring an animal’s bacterial load shortly after death. Though studies have shown that the genes involved in infection tolerance somewhat overlap with those involved in resistance, this has yet to be comprehensively determined for different pathogens and across genetically diverse lines (Ayres et al., 2008; Ayres and Schneider, 2008; Schneider and Ayres, 2008; Troha et al., 2018).

*D. melanogaster* lines also display variability in infection response within genetically identical individuals. This variation has been indirectly linked to stochastic differences in bacterial growth within the colonized host and variation in the onset of the animal’s immune response (Duneau et al., 2017; Ellner et al., 2021). The exact source of stochasticity has yet to be directly observed, largely due to a lack of tools capable of providing information on bacterial load in individual animals over time. Characterization of infection progression in flies has typically relied on destructive methods to establish bacterial loads at static time points, e.g. dilution plating. Dilution plating involves sacrificing individuals and quantifying bacterial load using colony counts from serially diluted fly homogenates. While these “snapshot” analyses can provide biological insight into mechanisms of infection progression and clearance, they are unable to generate longitudinal measurements of infections in individual animals (Chambers et al., 2019; Kutzer and Armitage, 2016). Dilution plating is also labor-intensive, as flies must be homogenized and plated one at a time. Due to the labor-intensive nature of the experiment, fine temporal measurement of immune dynamics over long time periods has historically been difficult. New strategies that enable high throughput, non-invasive measurement of pathogen load are needed to understand infection dynamics and outcomes of genetically similar and diverse populations.

Historically, a “go-to” method for non-invasive imaging in rodent models is bioluminescence (Love and Prescher, 2020; Zambito et al., 2021). Bioluminescence employs luciferase enzymes that oxidize luciferin substrates, producing photons of light (Kaskova et al., 2016). These photons can be detected through tissues in whole organisms, enabling sensitive and non-invasive readouts (James & Gambhir, 2012). Because no external excitation source is necessary, background signals are very low compared to other optical (e.g., fluorescent) readouts (Contag and Bachmann, 2002). Despite these advantages, bioluminescence has only been sporadically used in *D. melanogaster* (Brandes et al., 1996; Stanewsky et al., 1997; Stempfl et al., 2002). One potential reason for its limited use is that uniform delivery of the luciferin substrate by feeding is difficult, as feeding patterns can vary from fly to fly (Ja et al., 2007). One solution is to use autobioluminescent systems, which produce light without the need for exogenous substrate delivery. There are operons that produce both the luciferase and luciferin, allowing transgenic organisms to continuously glow (Kaskova et al., 2016). One popular autobioluminescent system derived from bacteria is the *ilux* system (Gregor et al., 2018). Engineered from the bacterial *lux* system, *ilux* emits blue light (490 nm) and exhibits enhanced brightness and thermal stability. While autobioluminescent systems have been used for decades to illuminate the spread of various pathogens *in vivo* (Cronin et al., 2012; Massey et al., 2011; Morrissey et al., 2013), they have yet to be applied for studying bacterial clearance in *D. melanogaster*.

Here, we report a novel method employing the *ilux* system for longitudinally monitoring bacterial load in *D. melanogaster*. By expressing the *ilux* operon in requisite bacteria and using photon count as a reporter for relative microbial load, we can non-invasively monitor infection progression and clearance over time in individual flies. Measurements can be performed in high throughput and with fine time resolution by housing the flies in 96-well plates and using automated image acquisition programs. With this method, we are able to observe distinct infection dynamics between lines of *D. melanogaster* as well as between genetically identical individuals. We anticipate this method will allow for more facile phenotyping of the immune resistance and tolerance in naturally varying lines and the observation of stochastic events, particularly late in infection, which are technically difficult to measure with current methods.

## Results

*Drosophila* infection models have historically required animal sacrifice to determine pathogen load, which limits measurements of each fly to one time point (Figure 1A). While this method has provided key insights into fly innate immunity and disease progression (Chambers et al., 2019; Chambers Moria et al., 2014; Duneau et al., 2017; Kutzer and Armitage, 2016; Lazzaro et al., 2006)), the fine dynamics of infection progression and tolerance among individuals remains difficult to measure in high-throughput. To address these limitations, we designed a non-invasive imaging strategy to monitor pathogen load over time (Figure 1B). We employed bioluminescent *E. coli* that constitutively expressed the *ilux* reporter (*ilux-Ecoli*) as a proof-of-concept platform (Gregor et al., 2018). The load of these autoluminescent bacteria could then be tracked post-injection, with photon flux reporting on relative pathogen count. Thus, we hypothesized that this method could be used to track differences in immune response between genetically distinct individuals and identify stochastic differences in infection progression within groups of genetically identical individuals.

**Figure 1.**
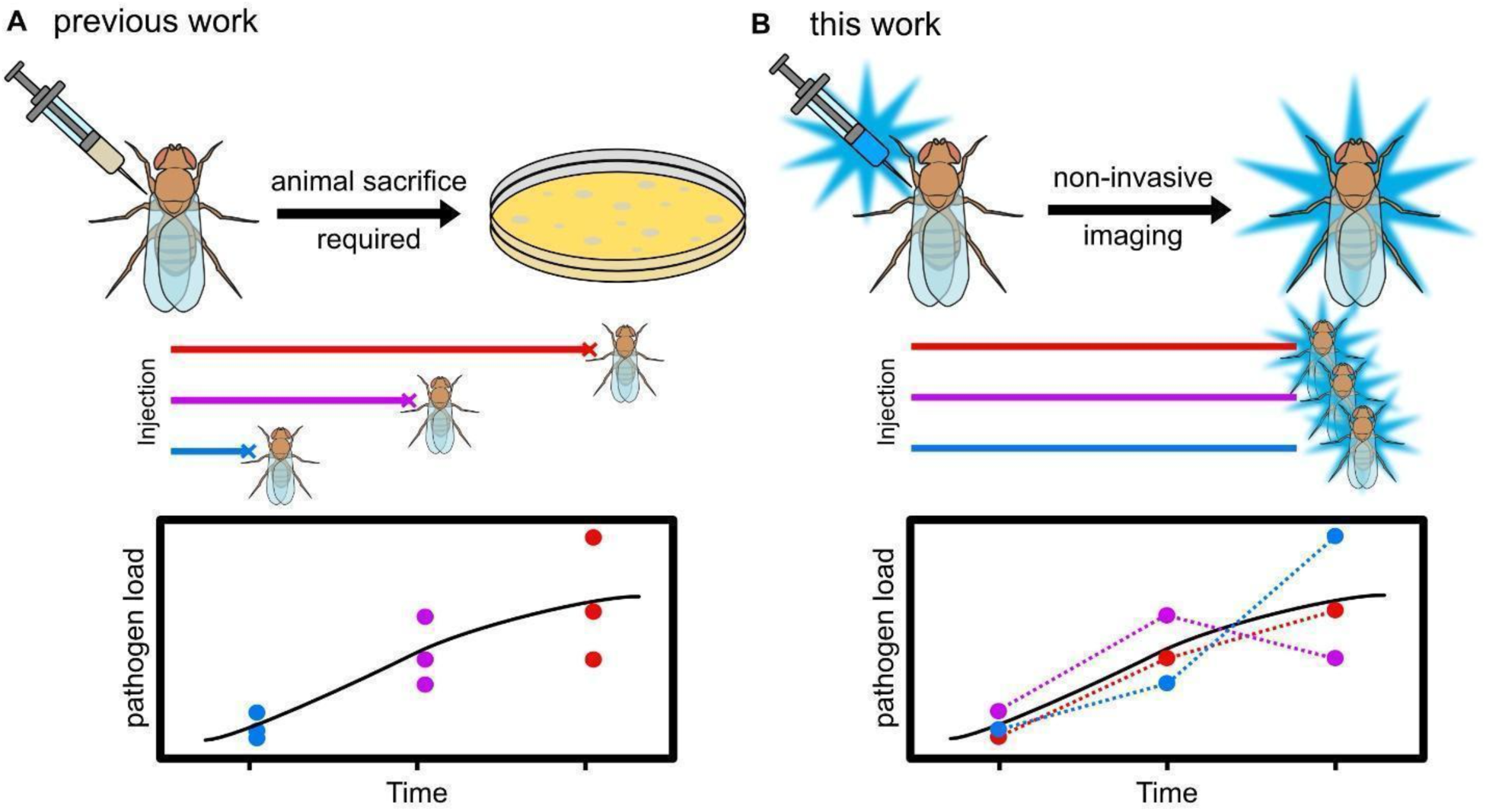
A novel method for non-invasive tracking of pathogen load over time. A) Previous methods for determining pathogen load require animal sacrifice. Larger cohorts are required for experiments as several flies must be sacrificed at the desired time points to check infection progression and clearance. B) This work presents a novel, non-invasive method to track pathogen load over time using bioluminescence. Thus, all flies can be individually monitored over time, allowing for a more comprehensive view of immune response.

To employ bioluminscence as a reporter for pathogen count, we first measured how photon flux correlated with bacterial optical density (OD) in liquid culture (Figure 2A). To do so, we serially diluted *ilux*-*Ecoli* in liquid culture and measured bioluminescent output. Photon counts correlated exponentially with bacterial OD. Since OD corresponds to bacterial colony forming units (CFUs) (Figure S1), relative pathogen abundances can be estimated from these photon measurements at this time point.

**Figure 2.**
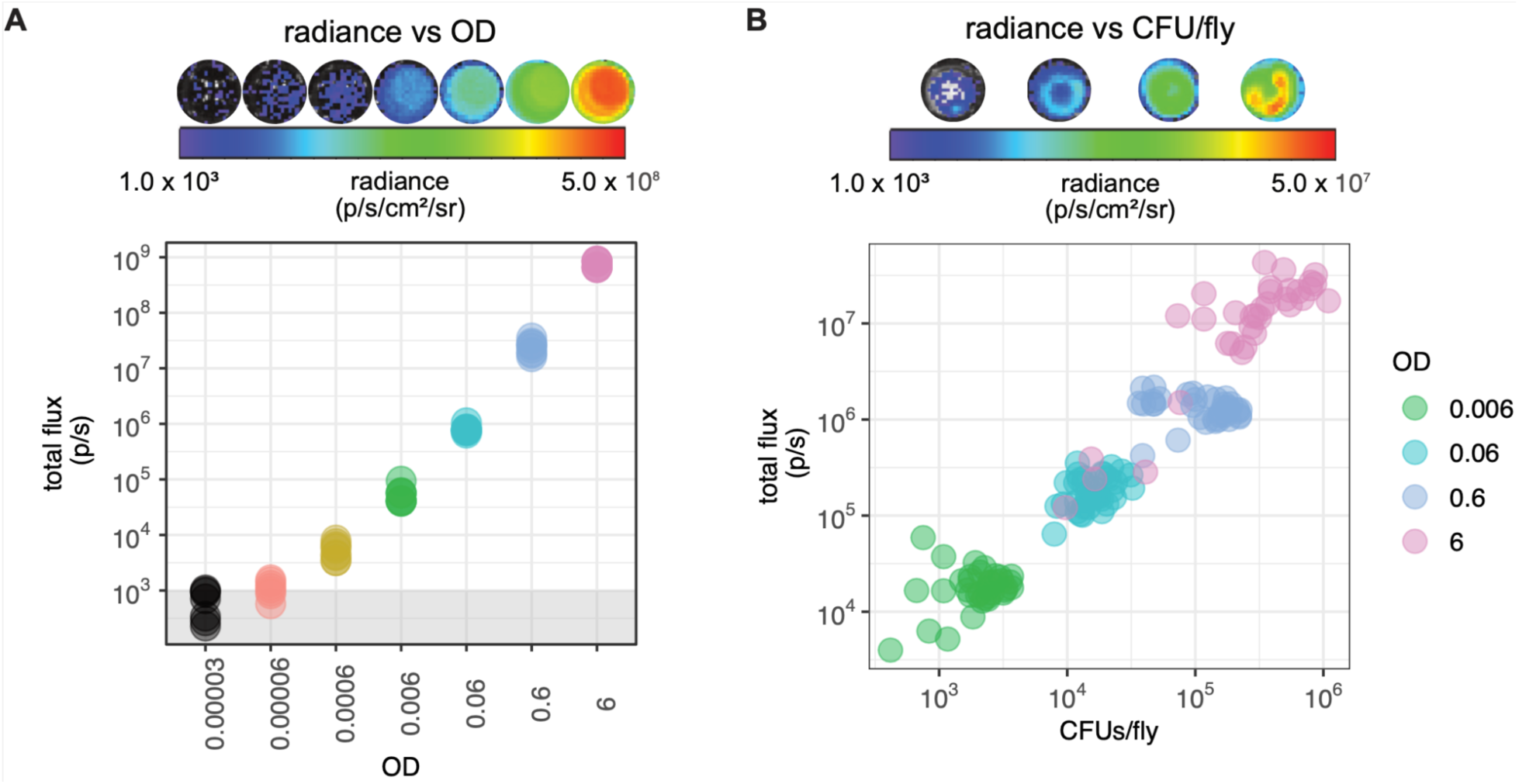
Photon flux is monotonically related to bacterial concentration. A) *ilux-Ecoli* were serially diluted in liquid cultures and assayed for photon output. Higher concentrations of bacteria yield higher photon fluxes. Plot shows eight measurements per OD (four biological replicates across two experiments). Well images are representative of the biological replicates. B) Wild-type flies were injected with different concentrations of *ilux-Ecoli* and assayed for light emission. A monotonic relationship was observed between radiance and CFUs injected. Graph shows data of 36 biological replicates per infection concentration, across two experiments. Well images above each graph are representative of images of the injected flies. The color scale is in units of photons/second/cm^2^/steradian. In both experiments, radiance was summed over the entire well to yield flux (photons/second) using the Living Image software.

We then sought to determine how flux correlates with bacterial count when injected into living flies by comparing the flux measurements of infected flies to CFU measurements generated by the current gold standard method, dilution plating. The *ilux* system emits blue light, with an emission maximum of 490 nm (Figure S2). While blue light is difficult to detect in the thicker tissues of mammalian model organisms, the fly cuticle is thin, and thus we hypothesized that the blue emission would be readily detectable in infected flies. Indeed, upon injection and imaging *ilux*-*Ecoli* in wild-type, male Oregon-R flies, we were able to reliably detect as few as 1000 CFUs. Given our injection volume of 34 nL, we estimated (Figure S1) and confirmed via dilution plating, that the flies injected with a solution of OD = 6, received approximately 100,000 CFUs. The thermal noise on the imaging instrument used (IVIS Lumina II) is between 10^2^ and 10^3^ photons/sec. Thus, we could not reliably image <200 CFU. A monotonic relationship was observed between CFU and photon flux indicating that flux can report on relative microbial count *in vivo* (Figure 2B). Notably, at the highest concentration of bacteria injected, there was a considerable amount of variation in initial dose, likely due to the injection needle clogging. The photon flux measurements for these animals showed a concordance with the CFU measurements, suggesting that this method can be used to filter out animals that received inconsistent doses.

To determine whether sexually dimorphic pigmentation of the cuticle affected this relationship, we also compared the best fit line of radiance/CFU of male and female flies. We found no differences between sexes (Figure S3). Thus, we were confident bioluminescence could be employed to determine pathogen count in both male and female flies. These results suggest that the *ilux-Ecoli* are sufficiently bright to be imaged through the cuticle, and that we can use standard curves of flies injected with known amounts of bacteria to calculate relative load at a given point in time. While flies bearing more pigmented cuticles (e.g. ebony flies) may attenuate signal more substantially, comparisons between individuals of the same cuticle hue would normalize such effects.

To determine whether bioluminescence could be employed for longitudinal tracking of pathogen load over time, we monitored flies for infection progression over several days. Given the duration of the experiment, the flies required housing both compatible with imaging and including food to prevent starvation. To this end, flies were housed in black 96-well plates by preparing small aliquots of food for each well and placing a glass sheet overtop of the plate (Figure S4). The glass sheet was secured with black electrical tape to mitigate aberrant photon scattering. With the housing in hand, we infected 48 wild-type flies with increasing concentrations of *ilux-Ecoli* and transferred them to individual wells in the prepared housing.

The flies were then imaged for four days, and photon fluxes recorded (Figure 3A-B). Since the flies were freely moving during the 3–5-minute image acquisition, they created a donut-shaped signal by traveling around the sides of the well (Figure 3A). We can account for fly activity in quantification by summing the photon count of the entire well. While the relative orientation of each fly in their respective wells could affect photon output, we anticipate these differences are averaged out by the length of the integration time and the similarity in movement observed for awake, freely moving animals.

**Figure 3.**
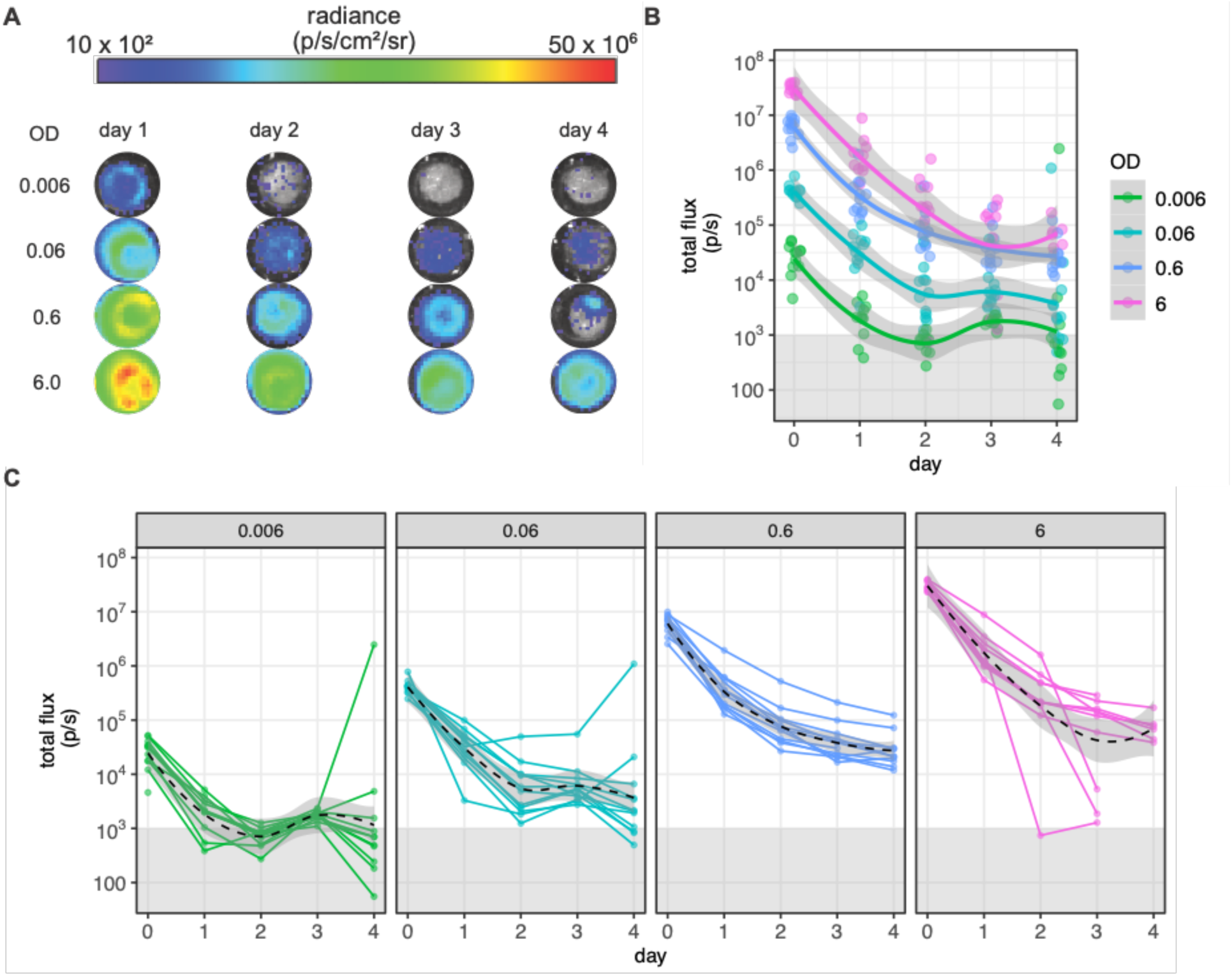
Bioluminescence can be used to track changes in pathogen load over time. A) Representative well images for different concentrations of bacteria injected in wild-type flies over four days. Higher initial concentrations were cleared to low, but detectable concentrations. Low initial concentrations were cleared below the limit of detection for the imaging instrument. Well images are representative of 48 injected individuals, 12 biological replicates per infection concentration. B) Average clearance patterns for different concentrations of *ilux-Ecoli* injected in individual flies over time. Photon counts were summed over the entire well where flies resided. The gray box shows the limit of detection of the imaging instrument. Solid colored line represents the average of the cohort. The gray band represents the standard deviation over replicates, dots represent one individual. C) Individuals display varied routes toward infection clearance, suggesting stochasticity plays a role in infection dynamics. Black dotted line represents the average, and the gray band represents the standard deviation. Gray box shows the limit of detection of the imaging instrument. Solid lines represent routes individuals took toward infection clearance.

Since wild-type flies mount a robust immune response against *E. coli* infection, none of the flies died during the course of the experiment. For the lowest doses of bacteria administered, photon flux values indicated that the infection had been cleared to levels below the detection limit. At higher doses, detectable amounts of bacteria remained on day 4, and the level of bacteria at day 4 is positively correlated with the initial load, consistent with previous reports (Chambers et al., 2019; Duneau et al., 2017). Further monitoring would reveal how long it takes for each fly to reach a steady state “set point” of bacterial load associated with a chronic infection state, whether there are fluctuations in this set point as a function of time, and the proportion of flies that may completely eliminate the infection.

Clearance patterns varied among flies receiving the same dose of pathogen (Figure 3C). For example, when looking at the trajectories from the two lowest doses, there are individual flies that experience resurgent infections. With dilution plating assays, it is impossible to verify whether a fly with a higher-than-average bacterial load at a later time point experienced a resurgent infection or failed to control the infection initially. Longitudinal measurements can distinguish between these scenarios. Some flies receiving the highest dose show markedly faster clearance dynamics than others.

To ensure the decreasing photon counts over time in wild-type flies were due to infection clearance rather than loss of the *ilux* plasmid, we repeated the same time course and imaged colonies from plated fly homogenates (Figure S6, S7). We observed minimal plasmid loss. We also found that feeding the flies ampicillin (the selection marker present on the *ilux* plasmid) had no effect on the proportion of luminescent vs non-luminescent colonies. We also investigated whether loss of the *ilux* plasmid was more pronounced in an actively proliferating infection. To this end, we infected immunodeficient imd^10191^ flies with *ilux-E*.*coli* and provided them with either normal fly food or food supplemented with ampicillin (Pham et al., 2007). When we compared the flux output, we found no differences between flies supplied with or without a selective compound (Figure S5). Thus, variations in flux observed in Figure 3 are reporting on true variation in bacterial load rather than variation due to loss of plasmid. Collectively, these results highlight how bacterial bioluminescence can be used to interrogate stochastic responses to infection in individual flies.

Historically, stochasticity of immune response in the first 12 hours of infection has been laborious to measure due to the need for fly sacrifice to obtain pathogen load information. We hypothesized our bioluminescent imaging strategy would be useful for studying differences in pathogen clearance during these critical first hours of infection. To test this capability, we used two fly lines, Oregon-R wild-type flies, and immunodeficient imd^10191^ flies. Wild-type flies are resilient to *E. coli* infection and clear this Gram-negative microbe easily, as shown in Figure 3. Imd^10191^ flies, by contrast, succumb to *E. coli* infection. These organisms bear a frameshift mutation in the IMD protein, effectively eliminating the immune response to Gram-negative bacteria. Therefore, we anticipated imd^10191^ flies would have different pathogen loads compared to wild-type files. Indeed, upon injection of a large quantity (approximately 100,000 CFUs) of *ilux-Ecoli*, imd^10191^ flies sustained high pathogen levels over time, while Oregon-R flies steadily cleared the infection as evidenced by reduced emission levels (Figure 4B). When handling highly concentrated pathogens for injection, variance in initial dose can occur. In this experiment, the imd^10191^ flies received a slightly lower initial dose of bacteria than the wild-type flies. However, by the two-hour time point, the imd^10191^ flies carried a higher load than the wild-type flies. Although we aim to deliver consistent initial doses, this result highlights a feature of this method: we can censor individual animals that receive aberrant initial doses. This quality control step is not possible with dilution plating-based methods.

**Figure 4:**
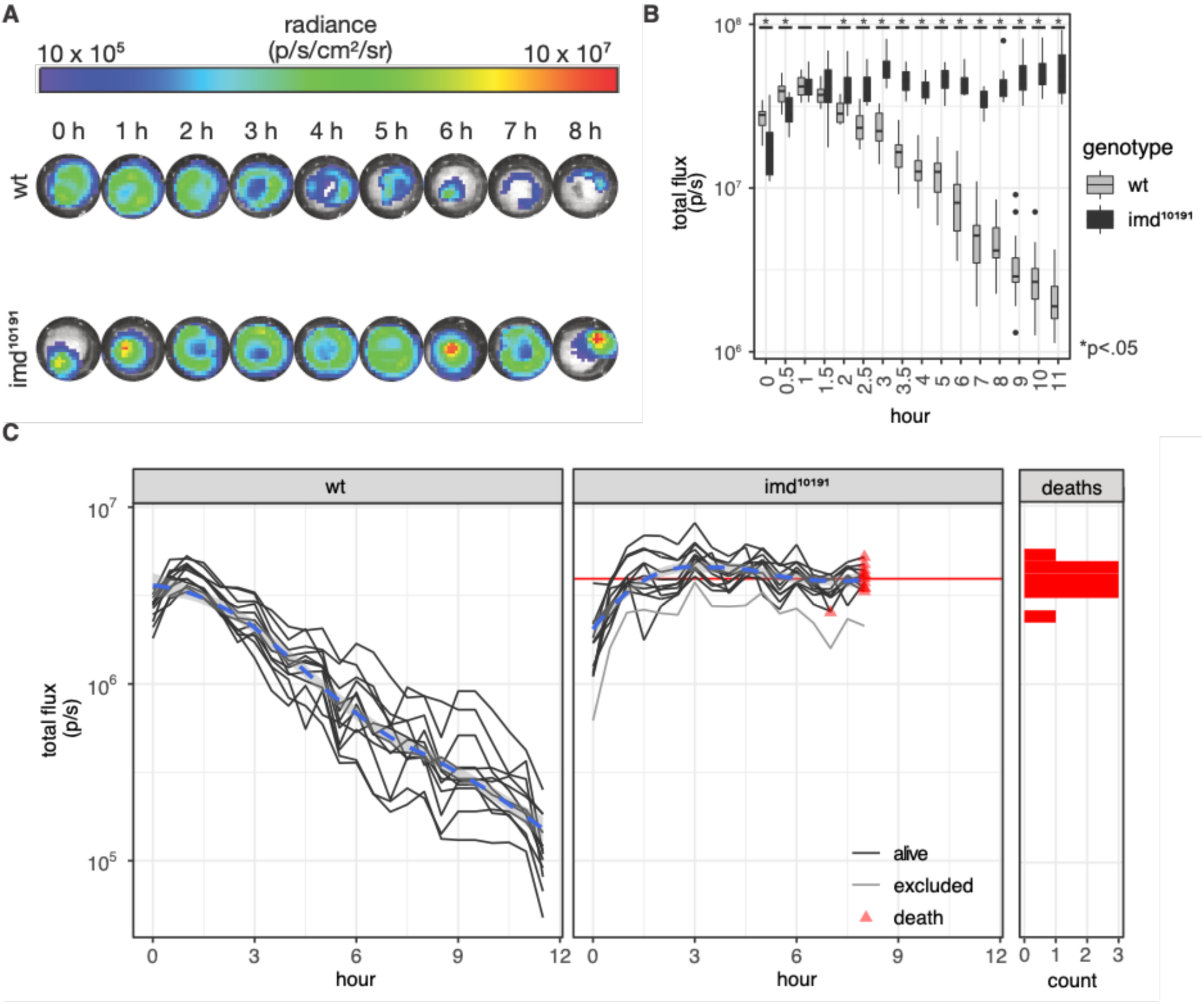
Longitudinal tracking of individual flies allows for deconvolution of community dynamics. A) Representative images of radiance measurements for wild-type (wt) and imd^10191^ injected with 0.034 µl of OD=6 *ilux*-*Ecoli*. Images show the first 8 hours, after which most imd^10191^ flies perished. Images are representative of 24 injected individuals, 12 biological replicates per genotype. B) Comparison of population level integrated total flux. Time points showing difference in mean between the two lines are demarked with an asterisk. Immunodeficient lines received a lower dose of infection than wt flies, but within an order of magnitude difference. This significance was lost by hour 1, with imd^10191^ bacterial load surpassing that of wild-type by hour 2. Data comprise 12 injected individuals. C) Comparison of wt and imd^10191^ individual tracks. Individual variation of immune response and pathogen clearance was observed in living flies (solid lines). Deaths are marked by red triangles, and the lines end. The blue dotted line shows the average of the cohort, while dark gray lines show the individual paths toward clearance or death. Death histogram shows the pathogen load upon death as estimated by total integrated flux. The red line on the imd^10191^ graph also demarcates the average of these values. The light gray line indicates a fly that received a lower-than-average initial dose and that was filtered out in the subsequent analysis. Data shown comprise 12 injected individuals.

Beyond quality control, we can actually use variation in initial dose to answer biological questions. For example, it has been shown that, within a genotype, varying initial dose by orders of magnitude yields differences in bacterial load of chronic infections. However, previous work relies on group averages and is unable to assess the impact of more subtle variations in initial infection load (Chambers et al., 2019). To address this question, we plotted the infection dynamics of individual flies (Figure 4C). To determine if the initial load of infection contributed to the differences in bacterial load at the end of the time course, we tested for correlation between flux at the time of injection and flux at the end of the time course for wild-type flies. We found the initial dose of pathogen does not significantly correlate with differences in the final load (Oregon-R R=0.19, Pearson’s product-moment correlation, Figure S8A), suggesting that smaller variations in initial infection dose do not determine the load at this timescale.

Previous work also suggests that bacterial load upon death is independent of initial load (Duneau, et. al 2017). In line with this work, we found the initial dose of pathogen did not significantly correlate with differences in the final load upon death in the imd^10191^ flies (R=0.24, Pearson’s product-moment correlation, Figure S8B). In the imd^10191^ line, we also looked to see if any correlation existed between the initial load of injection and the time it takes a fly to die after crossing the mean bacterial load upon death. We did not find a significant correlation between these two variables (imd^10191^ R=0.56, Pearson’s product-moment correlation, Figure S8C). The bioluminescent method thus reports on initial inoculation differences and can provide insight into alternative hypotheses for variance in infection dynamics among genetically identical populations.

In the experiment above, the high initial dose of bacteria killed the immune-deficient flies in less than 9 hours, which made it difficult to assess potential drivers of death. We posited that a lower dose of bacteria would kill immunodeficient flies more slowly and with greater variation in the time to death. To test this hypothesis, we injected wild-type and imd^10191^ flies with OD = 0.06 *ilux*-*Ecoli*. We then imaged flies every hour for 2 days post-infection. While both strains of flies were injected with the same concentration of bacteria, within the first three hours, pathogen load differed between the two populations (Figure 5A-B). Over time, the wild-type flies cleared the infection to low pathogen load and survived to the end of the experiment. Conversely, the imd^10191^ flies showed a gradual increase in pathogen load over time, with all flies succumbing to the infection by the end of the experiment. While initial loads varied between individuals, inoculation load again did not correlate with final bacterial load upon death for imd^10191^ individuals, or upon end of experiment for wild type individuals (Figure S9). This suggests that variance in initial load does not fully explain variation in bacterial load over time. In line with our hypotheses, we did observe substantial variance in time to death among the imd^10191^ individuals (Figure 5C).

**Figure 5:**
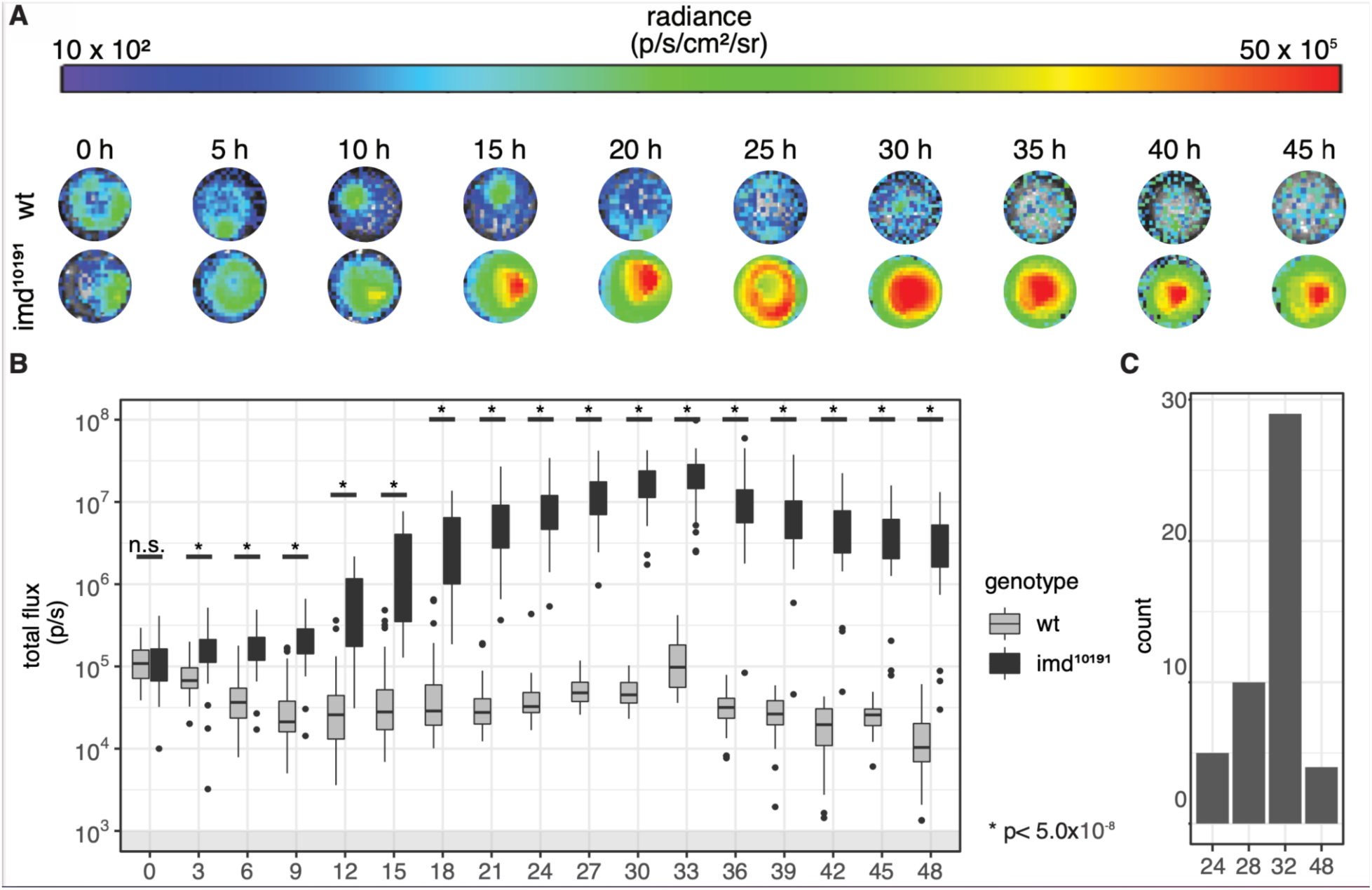
Individual infection tracking of immune-deficient flies allows for deconvolution of infection variation. Both wild-type (wt) and imd^10191^ flies were injected with 0.034µL of OD=0.06 *ilux-Ecoli* (n=48 for each genotype) A) Representative images of radiance measurements for wt and imd^10191^ flies injected with *ilux-Ecoli*. Images are representative for 96 individuals (48 biological replicates per genotype) and are shown in 5-hour intervals for the first 45 hours of infection. B) Summary of integrated total flux values for the wt and imd^10191^ in 5-hour intervals for the first 45 hours. Timepoints showing difference in mean between the two lines are demarked with an asterisk. By hour 5, both lines show differences in the ability to fight off infection with wild-type flies observed clearing the infection and imd^10191^ flies much higher bacterial loads. C) Histogram displaying time of death statistics for imd^10191^ flies. The majority of imd^10191^ flies died at hour 32, when integrated total flux reached its highest peak. Death data were compiled from 48 imd^10191^ individuals.

To further explore the variation in the infection progression in immunodeficient flies, we plotted their individual dynamics (Figure 6A). To identify groups of individuals showing distinct infection profiles, samples were hierarchically clustered using Euclidean dissimilarity (Figure 6B, Figure S10) (Montero & Vilar, 2014). This clustering requires that each individual have a measurement from all time points; therefore, we included bacterial load data post-mortem for flies that died prior to the end of the experiment. We separated the trajectories into four clusters, with the majority of flies falling into cluster 2 (blue, n=29) or cluster 3 (yellow, n=13), and clusters 1 and 4 having three flies each (magenta and green, respectively). Clusters 2 and 3 appear to separate based on the lag time to unchecked bacterial growth, with cluster 2 having a lag time of 5-10 hours and cluster 3 having a lag time of 10-15 hours after infection. Clusters 1 and 4 appear to cluster primarily based on receiving a below-average initial dose (cluster 4) or on bacterial dynamics post-mortem (cluster 1). No cluster appeared to correlate with time of death (Welch two sample t-test on every cluster combination, Figure S11). While all samples ultimately succumbed to the infection, these distinct dynamics -- particularly the difference between bacterial lag time and growth rate variation -- would have been missed using previously established methods.

**Figure 6:**
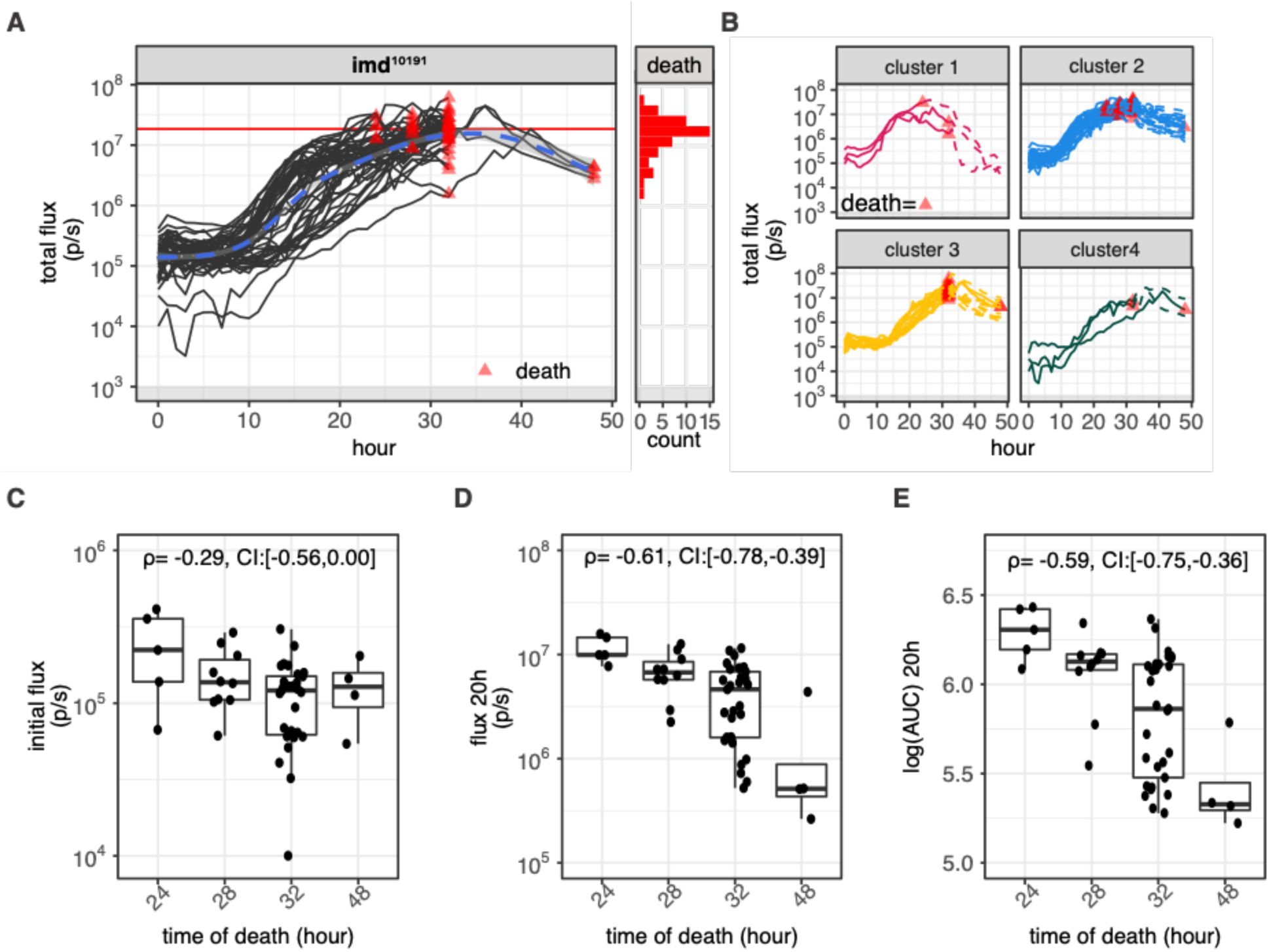
Individual infection tracking of oregon-R flies shows two distinct pathways towards bacterial clearance. A) Individual tracks of infection in live imd^10191^ flies (black lines, n=48); deaths are marked as a red triangle and the end of the line. All flies died by hour 48 and the mean radiance upon death was 18.6×10^6^ (solid red line). Threshold of accurate detection is demarcated with a gray box. Histogram shows the distribution of integrated photon flux (serving as a proxy for pathogen count) of imd^10191^ flies upon death. B) Individual tracks of infection separated by clusters. Clusters were assigned via hierarchical clustering using Euclidean dissimilarity. Four distinct groups were assigned with cluster 2 and 3 containing the majority of samples (cluster 2= 29, cluster 3=13) and clusters 1 and 4 containing 3 samples each. Data are for 48 imd^10191^ individuals. C) Spearman rank correlation (ρ) and the 95% bootstrapped confidence interval (CI) between initial flux and time of death for imd^10191^ flies. D) Spearman rank correlation and CI between total flux at hour 20 and time of death. E) Spearman rank correlation and CI between the log transformed area under the curve (AUC) up until hour 20 and the time of death. The curve here refers to the flux (p/s) vs time curve. In all cases CI of Spearman correlation coefficients were computed by bootstrapping 10,000 synthetic datasets and computing the correlations on these datasets. 2.5 and 97.5 percentile values of the sampled correlations are reported.

We further investigated potential sources of the variability observed in time to death of the imd^10191^ flies. Several possibilities exist to explain these differences in dynamics, including variation in the injury upon infection, variation in the initial pathogen load, or differences in the physiological state of the fly. To examine if this variance in time to death can be explained by differences in the initial load of infection, we performed Spearman rank correlations between time of death and the initial load. We also measured the correlations between time of death and load at 20 h post injection and the area under curve (AUC) of pathogen load at 20 h, as this is prior to any of the animals dying. We found the initial load was negatively correlated with time of death (ρ = −0.29, 95% bootstrapped confidence interval = (−0.56, 0.00)) (Figure 6C). Intuitively, this aligns with expectations: flies receiving higher inoculations will die more quickly. The strongest correlation was found between time to death and observed load at 20 h post injection (ρ = −0.61, 95% bootstrapped confidence interval = (−0.78, −0.39)) (Figure 6D). A similarly strong correlation was observed between time to death and area under the flux curve at 20 h post injection (ρ = −0.59, 95% bootstrapped credible interval =(−0.75, −0.36)) (Figure 6E). These correlations highlight how factors beyond initial load contribute to the time of death observed in individual flies.

Unlike the immunodeficient line, all of the wild-type flies survived well past two days, with the overall trend of clearing the infection (Figure 7A). Despite the average drop in pathogen load, we did observe large amounts of variance, particularly between hours 10 and 20 of infection. Hierarchical clustering sorted the individual traces into groups that corresponded to low, intermediate or high pathogen load in the first 25 hours of infection (Figure S12). In studying this data, we noticed that some individuals experienced a resurgence in infection, and these individuals fell into more than one cluster. Therefore, we highlighted profiles based on the presence of a secondary distinct increase of pathogen load after the initial infection (group 1 : magenta, n = 10) rather than a sustained gradual decrease (group 2 : blue, n = 38, Figure 7B). We found the initial dose of infection did not significantly correlate with the final flux between both groups (group 1 R=-0.41, group 2 R=-0.15, Pearson’s product-moment correlation, Figure S13A) and that neither the initial nor the final flux measurements were significantly different between these groups (group 1 vs. group 2 initial p=0.2, group 1 vs. group 2 final p=0.45, Welch two sample t-test, Figure S13B and C). Despite the presence of an increase in pathogen load in some flies, the population ultimately converges towards controlling the infection. We expect that this method’s ability to report fine differences in pathogen load over time will enable future research to uncover the origins of such stochasticity and better characterize the paths hosts may take to clear an infection.

**Figure 7:**
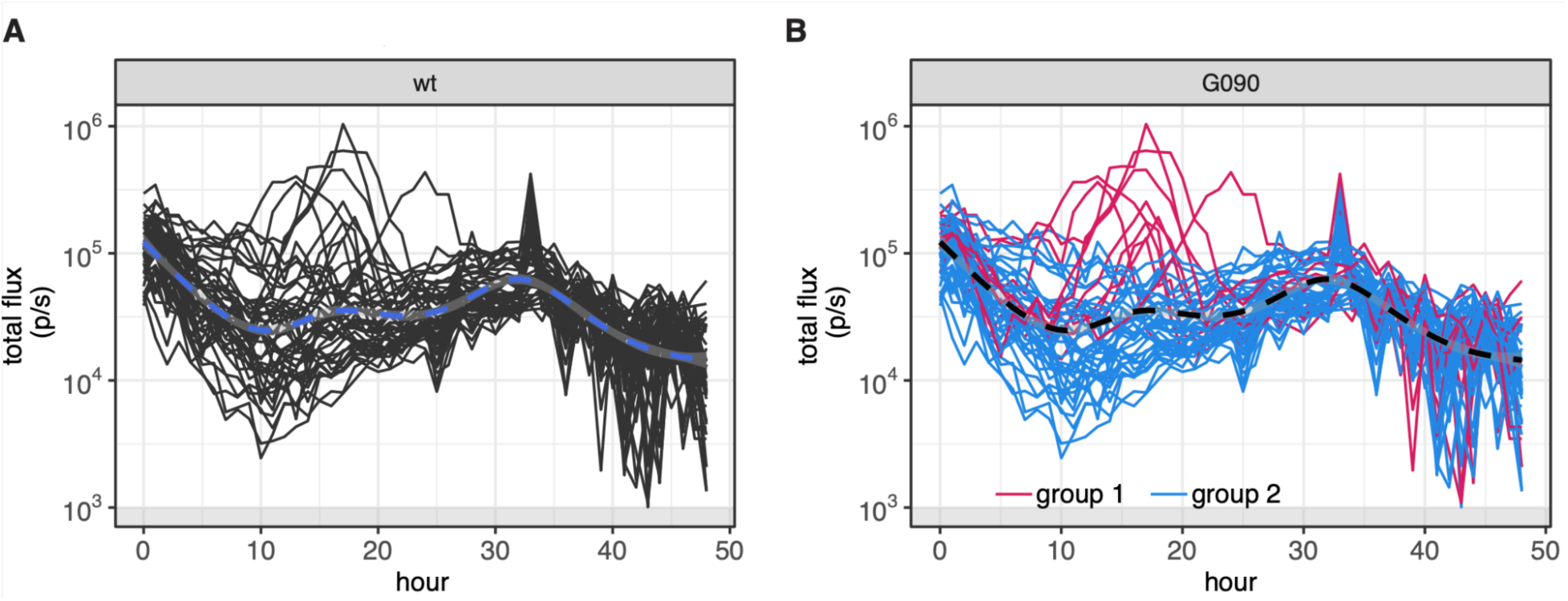
Individual infection tracking of wild-type (wt) flies shows two distinct pathways towards bacterial clearance. A) Individual tracks of infection in live flies (black lines). No flies died during this time course. Threshold of accurate detection demarcated with gray box. B) Individual tracks of infection are grouped by the presence of a secondary peak during the infection process. While all flies showed a decrease in bacterial load by the 48-hour mark, a subset of flies (n = 10) showed an increase in bacterial load between 10 - 25 hours (magenta lines). Dashed lines represent the mean trajectory for the genotype. The noise in some of the later time points is due to a shortening of the acquisition time for these animals, which were housed in the same plate as the highly luminescent immunodeficient animals shown in Figure 6.

## Discussion

Here we show that employing an autobioluminescent bacterial reporter enables simple, non-invasive pathogen load determination in *Drosophila melanogaster*. We show that total flux can report on relative changes in bacterial load over time. The non-invasive feature of this method fundamentally changes our view into infection dynamics by enabling longitudinal tracking of infections in individual animals. Traditional dilution plating approaches only allow for the measurement of an average infection trajectory in a population of genetically identical individuals, while longitudinal measurements allow us to reveal individual variation in dynamics and features such as resurgent infections. Further, these traces can be used to test hypotheses about what features (e.g. initial dose, time to immune system engagement, or colonization progress) drive the ultimate outcome of infection. The simple housing requirements and rapid acquisition times allow for the efficient measurement of large numbers of animals or genotypes at fine time resolution. Because each animal is repeatedly sampled throughout the experiment, the longitudinal measurements effectively reduce the sample size needed to identify differences in distinct genotypes of flies. The decrease in sample size, coupled with the increase in throughput, will make previously laborious genetic screens to identify new components of the immune response more accessible.

This approach may serve a complementary role to approaches integrating fluorescently labeled bacteria, which have been used to track the localization of some microbes during infection, particularly in larvae (Mansfield et al., 2003; Muniz et al., 2007; Rutschmann et al., 2002). However, fluorescent microbes have been difficult to use as a quantitative readout of bacterial load due to *Drosophila* autofluorescence and generally lower signal to noise. Furthermore, illumination with high powered light sources (i.e. UV lamps) can damage tissue and affect animal behavior (Du et al., 2016; Kim and Johnson, 2014; Milinkeviciute et al., 2012). Bioluminescence can thus offer a less perturbing alternative to fluorescent methods.

The flexibility and simplicity of this method should enable its use in a wide range of settings. For example, the *ilux* cassette can be manipulated via molecular cloning, allowing expression in a wide variety of pathogens. To establish the system in a new organism, a vector that contains a species-appropriate promoter, origin of replication, and resistance marker should be identified, and controls should be done as in Figures 2, S5, and S6 to ensure the relationship between light output and bacterial load can be used to measure relative changes in bacterial load over time. Depending on the microbe, features like toxin-antitoxin cassettes or chromosomal integrations can be used to improve the quantitative performance of the system (Hayes, 2003; Yamaguchi et al., 2011). We have also demonstrated that this system works well in multiple fly genotypes. Though certain fly genotypes may have an impact on light emission, e.g. the dark cuticle of *ebony* flies, this approach can still be used to compare relative bacterial loads within an individual over time or between individuals of the same genotype. The fly housing requires only simple components found in most laboratories. We also examined the feasibility of using a plate reader instead of an IVIS imager for luminescence measurements (Figure S14). Using flies injected with bacterial doses spanning 6 orders of magnitude, we found that while the IVIS has better sensitivity in detecting low pathogen loads, the plate reader performed comparably to the IVIS at higher doses. This indicates that the plate reader may be useful for experiments in which the bacterials loads are likely to be above the threshold of detection.

The longitudinal measurements and temporal resolution achieved using bioluminescence enables novel observations of infection dynamics. For example, although we found wild type flies cleared *ilux-Ecoli* in all experiments, some individuals experienced a resurgence in infection upon inoculation with moderate doses of bacteria (Figure 7). Using previous dilution plating methods, resurgence cannot be definitely identified. It would appear to be variation between sacrificed individuals at discrete time points. Using our bioluminescent method, resurgence can be easily visualized as an increase in total flux observed in certain individuals of a population and future studies may investigate the causes and predictors of a resurgent infection.

New insights were also gleaned from experiments with immunodeficient flies. We found a moderate correlation between the initial load and time to death for immunodeficient flies, and a stronger correlation between time to death and bacterial load at 20 h post injection for individuals receiving a low initial dose of *ilux-Ecoli* (Figure 6D). Therefore, it seems both variation in initial load and infection progression combine to determine the ultimate time of death. The variable paths flies take toward a fatal infection can now be investigated in fine detail using the temporal resolution afforded by our bioluminescent method. Ongoing work involving integration of additional luminescent reporters to label immune system components would enable even more thorough investigation into how different variables can contribute to infection outcome.

In summary, non-invasive tracking of pathogen load in *Drosophila melanogaster* over time offers many advantages when compared to traditional methods. Dilution plating requires animal sacrifice to determine average pathogen load at static time points, limiting the ability to investigate individual infection dynamics, particularly over long time scales. The bioluminescent method presented herein offers a facile approach to non-invasively monitor pathogen load on the individual level. Because images may be automatically acquired on the minute-time scale simultaneously in many flies, bioluminescence enables infection progression to be monitored with exceptional resolution across long time periods and in high throughput. This may enable previously-difficult genetic screens, for example, to examine variation in infection tolerance between naturally varying fly lines or dynamic-dependent phenotypes, like propensity to endure a resurgent infection. Integrating this method with other visual reporters of host state, e.g. the activation of immune signaling pathways, may further illuminate pathogen host-dynamics (Troha and Buchon, 2019). Using this method, we can begin to quantify the contributing factors that result in stochasticity, resurgence, and ultimate infection outcome.

## Acknowledgments

We would like to thank Dr. Larry Marsh and Dr. Neal Silverman for sharing their fly lines to make this work possible and Zi Yao for his assistance with the pilot experiments. We would also like to thank David Duneau for useful feedback on an earlier version of this manuscript. This work was supported by NSF/BIO/MCB grant 1953324 to ZW, a Convergence Accelerator Team award from NSF-Simons Center for Multiscale Cell Fate Research to JP and ZW, and a seed grant from the UCI Infectious Disease Science Initiative to JP and ZW.

## Materials and Methods

### Preparing *ilux E. coli*

An *E. coli* strain harboring a plasmid with the *ilux* operon (*ilux* pGEX(−)) was obtained from Addgene (plasmid # 107897, deposited by Stefan Hell) and streaked on an LB agar plate with ampicillin (100 µg/mL) to afford single colonies. A single colony was picked and grown in 5 mL LB broth containing ampicillin (100 µg/mL, LB-AMP). The culture was miniprepped according to the manufacturer’s instructions (kit purchased from Zymo Research). Plasmid (10 ng) was transformed into chemically competent TOP10 *E. coli* (20 µL). The transformant was recovered with SOC (50 µL) for 30 mins at 37 °C and 25 µL plated on an agar plate containing ampicillin. A single colony was picked and expanded in LB-AMP, and a glycerol stock was made for long term storage at −80 °C (500 µL culture with 500 µL 50% v/v glycerol). This glycerol stock is referred to as *ilux*-*Ecoli*.

### *Drosophila* lines and rearing

Oregon R and imd-10191 were used for this study (Pham et al., 2007). Both lines were reared on standard cornmeal media at 20°C (Brent and Oster 1974). Four-day old male and female flies were collected for injections to ensure full replacement of the larval fat body by the adult fat body (Johnson & Butterworth, 1985).

### *Drosophila* infection induction

Prior to infection, *ilux-Ecoli* was cultured in liquid LB-AMP on a shaker at 37°C for 8 hours. Bacteria were then pelleted using a table-top micro-centrifuge at 5000 rpm/g and resuspended in 200<u>µl</u> of 1X phosphate buffered saline. Optical density at 600 nm was then measured using a NanoDrop 2000 (Thermo Fisher). Injection solutions were prepared at the appropriate OD by dilution in additional PBS. Flies were injected with 34 nL of bacterial solution using Narishige IM 300 Microinjector along the scutescutullar suture and immediately placed into black 96-well plates (Grenier Bio One). For time course experiments, 96-well plates for imaging were prepared by punching out circles of standard cornmeal media and placing these at the bottom of wells before placing flies into the plate. A 4-inch by 6-inch glass cover was placed on the plate during the duration of the time course to prevent individual escape. For single time point measurements, flies were placed in 96-well plates lacking food.

### Dilution plating: *ilux-Ecoli*

To determine the concentration of *ilux-Ecoli* at per OD measurement, *ilux-Ecoli* was grown in 10 mL LB-AMP on a shaker at 37°C for 6-7 hours while the bacteria was still in exponential growth phase. Bacteria was then pelleted and resuspended in 200 uL 1X phosphate buffered saline and OD at 600 nm of the solution was determined using a NanoDrop 2000 (Thermo Fisher). A stock solution of OD 1 was prepared and serially diluted using 1x PBS by 6million fold. CFUs were then quantified in two ways. The first manner was using 5µl of solution for each dilution which was spot plated in triplicate on LB plates supplemented with 100µg/ml ampicillin. The second manner was using 90 µl of solution which was plated on LB plates supplemented with 100 µg/ml ampicillin. Colonies were then counted for each dilution step to determine the concentration of CFUs at OD1 for both methods (Supplementary Figure 1). To determine the relationship between *ilux-Ecoli* concentration and total flux, a solution of bacteria was prepared as described above, 90µl of each dilution was then placed into black 96 well plates for imaging (see “imaging parameters” section below).

### Dilution plating: infected *Drosophila*

To determine the concentration of bacteria injected into individual flies, flies were suspended in 250 µl of 1X phosphate buffered saline and homogenized (Krupp & Levine, 2010). Homogenate was then used for stepwise 5-fold serial dilutions. 5 µL of each dilution in the series was then spot plated in triplicate (3 technical replicates per biological replicate) on LB plates with 100 µg/ml ampicillin. Colonies were then counted for each dilution step and the mean of the technical replicates was used to determine the concentration of CFUs per fly. For plasmid loss determinations, homogenate and serial dilution was prepared in the manner described above. Next 90 µl of solution from each dilution in the series was plated on individual LB plates not supplemented with ampicillin. Plates containing colonies were then counted for each dilution step to determine the concentration of CFUs per fly and imaged for luminescence output as described in the “imaging parameters’’ section. Thus, emissive and non-emissive colonies could be distinguished to determine the proportion of colonies that had lost the *ilux* plasmid.

### Imaging parameters

All imaging analyses were performed in black 96-well plates (Grenier Bio One) prepared as described above, or on agar plates for plasmid loss studies. Individual experiments took place on seperate days, whereas individual infected flies were considered biological replicates. Plates containing flies were imaged immediately post-injection unless otherwise stated. Injected flies and agar plates containing *ilux-Ecoli* were imaged using an IVIS Lumina II (Xenogen) CCD camera chilled to −90 °C. The stage was kept at room temperature (25 °C) during the imaging session, and the camera was controlled using Living Image Software. The exposure time was 1 s - 10 min depending on the brightness of the sample, and the data binning levels were set to medium. Background thermal noise levels of the instrument were determined via acquisition of images containing no bioluminescent materials for all integration times used in this study. Maximum photon values observed in these “dark charge” images were used as the absolute background levels of the instrument. Because the dynamic range of the instrument is three orders of magnitude, relative background levels can change depending on the radiance observed from the brightest sample present. In these cases, relative background levels would be three orders of magnitude lower than the highest pixel radiance observed. Radiance was integrated over regions of interest and quantified to total flux values using the Living Image software. Raw luminescent images were analyzed using FIJI (Schindelin 2012). For plasmid loss studies, CFUs on plates were quantified by hand, luminescent colonies were quantified by importing for analysis and counting in FIJI.

### Estimate of detection limit

To estimate the threshold of detection of our assay we first measured the total flux from wells containing only fly media which was found to range between 10^2^ and 10^3^ photons/sec. To be conservative in our estimates we selected 10^3^ photons/sec as our threshold. We then took the model2 linear regression of total flux measured by the IVIS versus the CFUs/fly calculated via dilution plating. Finally we calculated our CFU/fly detection limit by inputting our flux detection limit and solving for CFUs.

## Data analysis

All luminescence data was imported into R 3.6.0 for analysis and visualization (R Core Team. 2019, Wickham H. 2016, Dowle M(2019), Wickham H. 2021, Wickham et al., 2019). Model2 regressions were performed using the package lmodel2 1.7-3 (Legendre 2018). Euclidean dissimilarity and hierarchical clustering were performed using package TSclust 1.3.1 (Montero & Vilar, 2014).

## Data availability

All flux data generated during this study as well as the code used to analyze the data and generate figures are available for download at github.com/WunderlichLab/ilux_infection_tracking.

## Notes

### Competing Interest Statement

The authors have declared no competing interest.

