## Supplementary Information for "Longitudinal monitoring of individual infection progression in *Drosophila melanogaster*"

**Supporting Material for "Longitudinal monitoring of individual infection progression in *Drosophila melanogaster*"**

**Figure S1: *ilux-Ecoli* is calculated to be ~550,000 CFUs/µl at OD=1**
**Figure S2: Spectroscopic properties of *ilux***
**Figure S3: Male and female flies show no differences in relationship of CFUs injected to radiance detected**
**Figure S4: Schematic of the plate set up used for housing and imaging flies**
**Figure S5: Media supplemented with ampicillin did not affect flux output over the course of infection**
**Figure S6: *ilux-Ecoli* injected into flies show minimal plasmid loss over time**
**Figure S7: Representative images of colonies used to measure plasmid loss**
**Figure S8: Initial dose of infection does not result in differences in final load or time to death in Oregon-R or imd$^{10191}$**
**Figure S9: Initial dose of infection does not result in differences in final load in imd$^{10191}$**
**Figure S10: Unsupervised clustering of imd$^{10191}$ results in 4 distinct clusters**
**Figure S11: Comparison of hour of death to assigned cluster**
**Figure S12: Unsupervised clustering of Oregon-R infection profiles does not separate trajectories by infection resurgence**
**Figure S13: Initial load of infection does not result in differences in final radiance or resurgence of infection**
**Figure S14: Autobioluminecent bacteria is detectable by both IVS and plate readers**

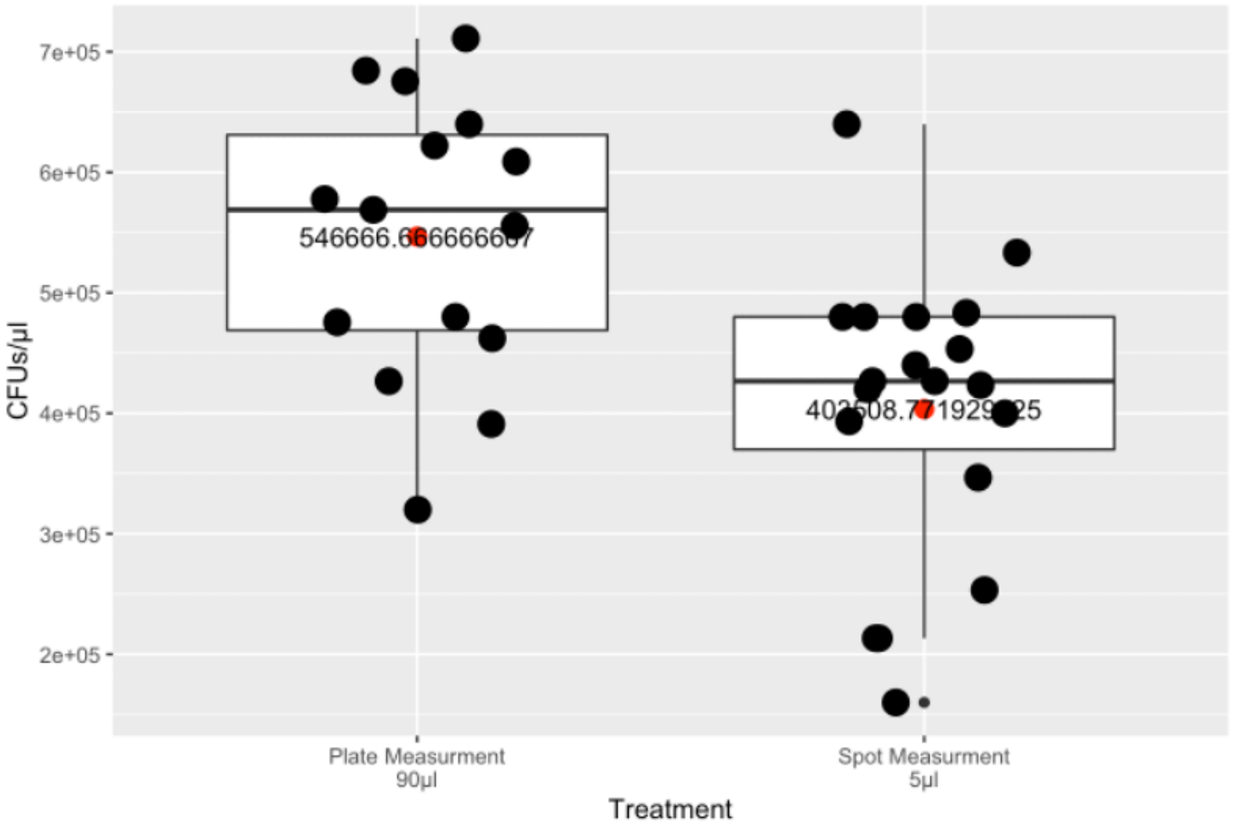


**Figure S1. *ilux-Ecoli* is calculated to be ~550,000 CFUs/µl at OD=1.** Calculated CFUs/µl for two dilution plating experiments. *ilux-Ecoli* solution were prepared, serially diluted and plated using one of two methods (see Methods). From each dilution, either a single 90µl aliquot was plated on a single plate (left) or six 5µl aliquots were plated on a single plate (right). Calculated CFUs per µl for both measurements were within an order of magnitude of each other. We chose the 90µl plate measurements for our final calculations given that the larger aliquots make them less likely to be skewed by stochastic sampling.

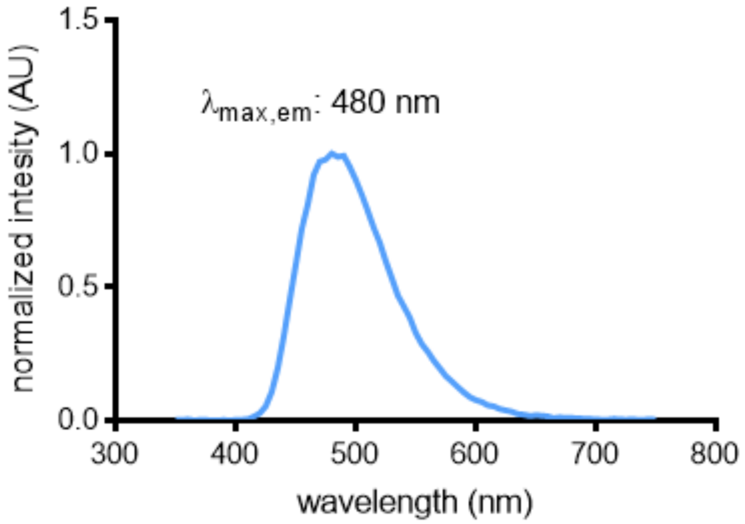


**Figure S2. Spectroscopic properties of *ilux*.** *ilux-Ecoli* were cultured in LB-AMP for 24 h at 37 ℃. An aliquot of cells (700 µL) was transferred to a quartz cuvette, and the bioluminescence spectra was recorded at 25 ℃. Luminescence values were normalized such that the maximum value equaled one and plotted in GraphPad Prism 5.

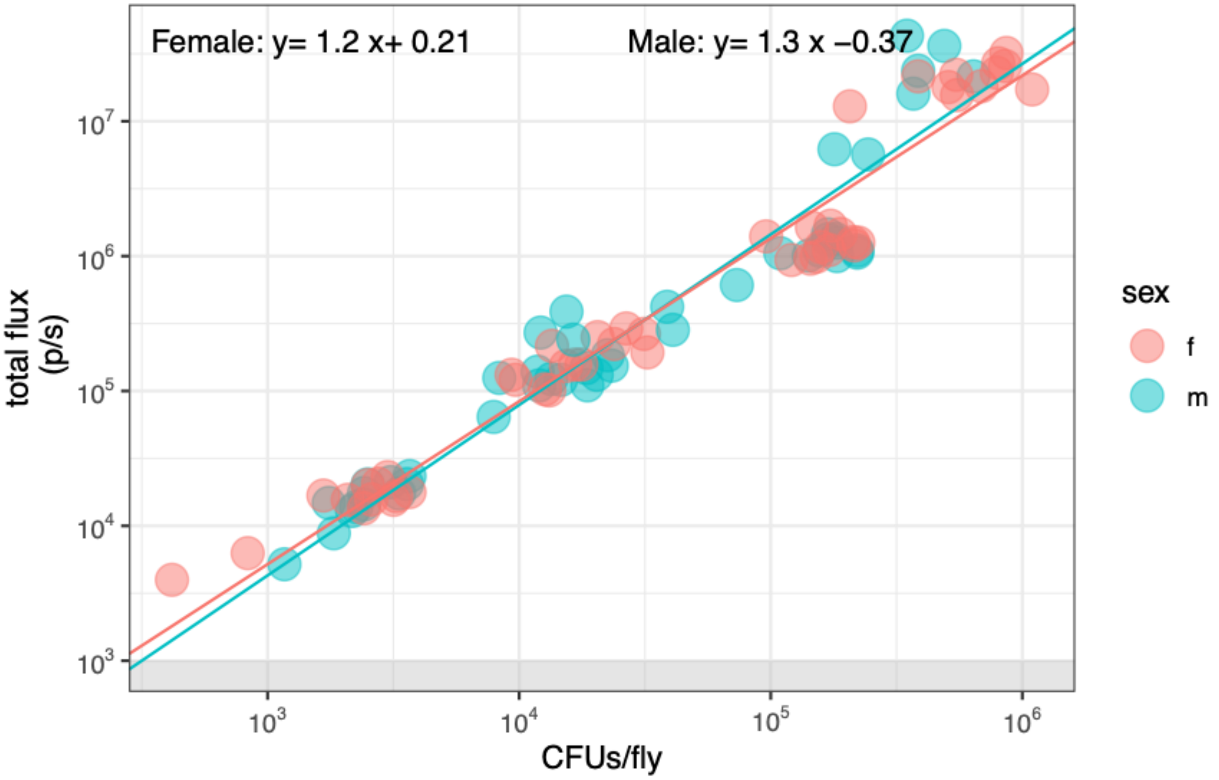


**Figure S3: Male and female flies show no differences in relationship of CFUs injected to radiance detected.** Male and female flies were injected with one of four doses (OD = 6, 0.6, 0.06 or 0.006) of *ilux-Ecoli*. Standard major axis regression of log-transformed data for each sex showed no significant differences between slope (*m*) of the lines male *m*= 1.3 (CI: [1.37,1.16]) female *m*=1.2 (CI:[1.28,1.14]). While there is a small difference in the intercept, we conclude that the relationship between photons/CFU is similar between female flies and the more pigmented male flies. See Methods for more details.

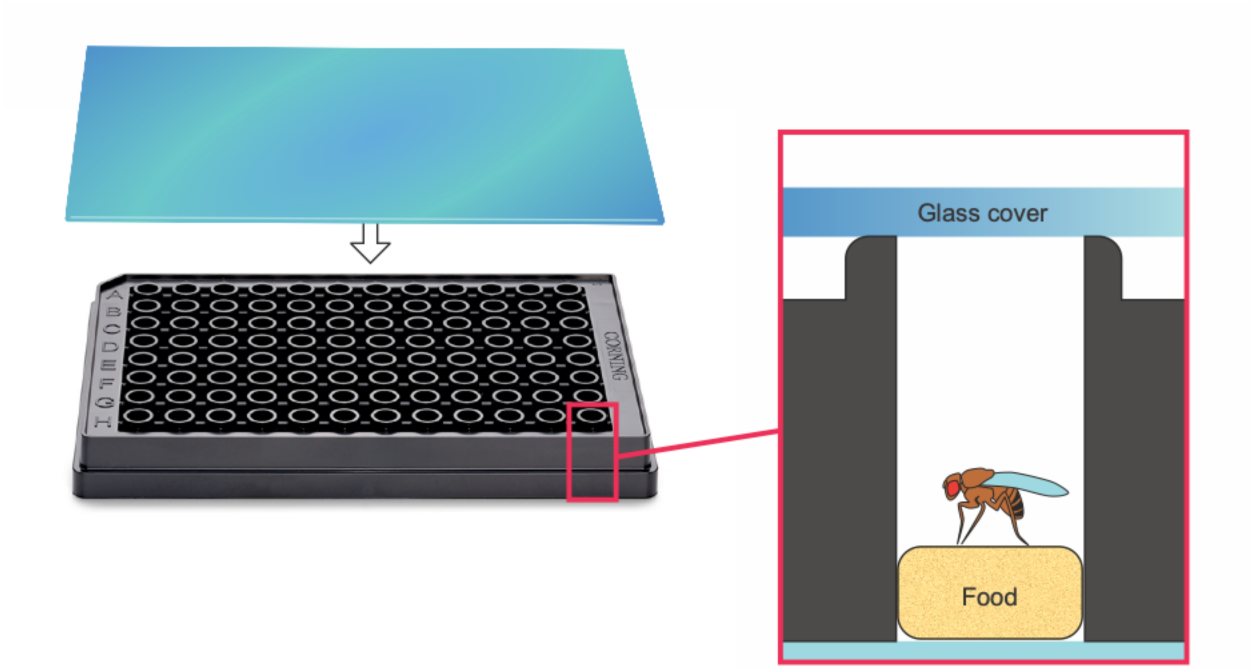


**Figure S4: Schematic of the plate set up used for housing and imaging flies.** Black-walled, clear-bottomed 96-well plates with a 2 mm thick glass cover were used to house flies for the duration of the imaging time course. For food, 2-5 mm thick disks of standard cornmeal media were placed at the bottom of each well before placing individual flies inside. Disks were prepared by using the end of a 5 mL serological pipette to "punch out" disks from solid media.

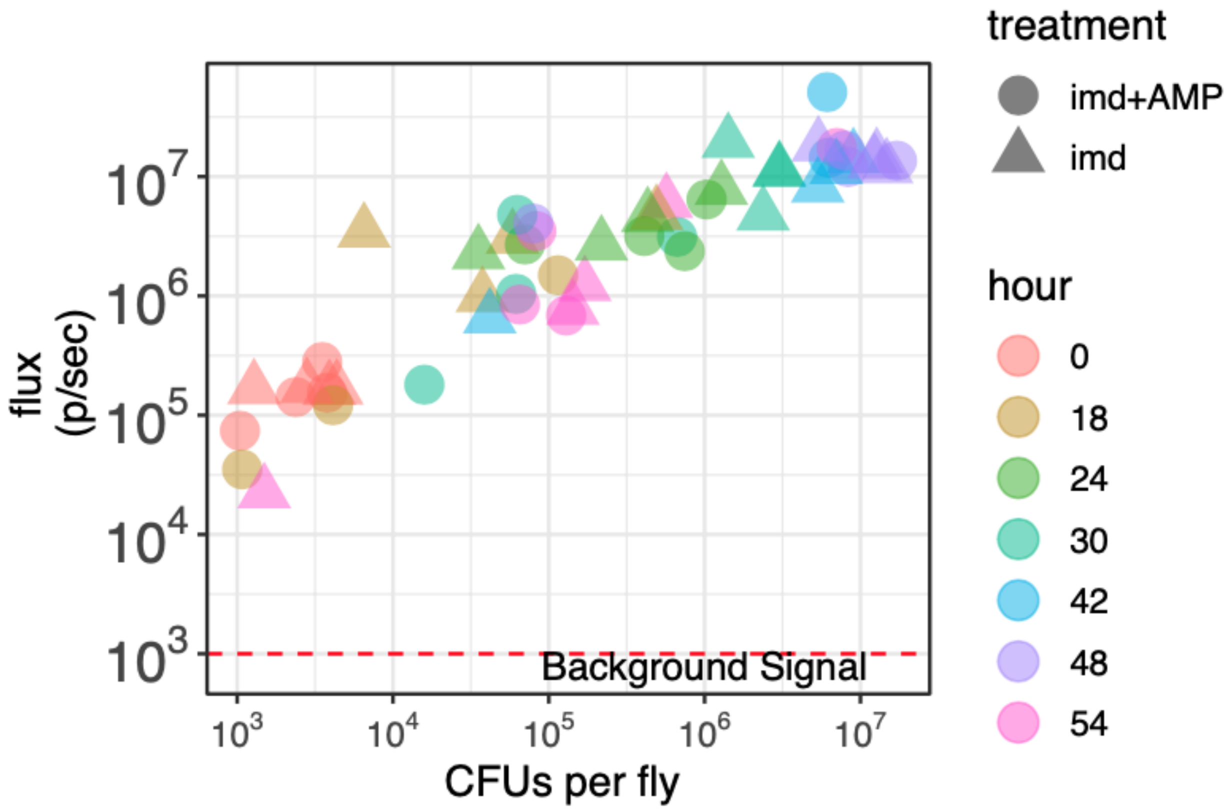


**Figure S5: Media supplemented with ampicillin did not affect flux output over the course of infection.** To determine if the absence of plasmid selection may affect flux output during the course of a proliferating infection, male imd$^{10191}$ flies were inoculated with an 34nl of OD of 0.06 ilux bacteria. Flies were housed in 96-well plates with standard cornmeal media or media supplemented with 100µM ampicillin to select for the ilux plasmid. At each time point the plates were imaged and a subset of flies were sacrificed for pathogen quantification via dilution plating. We found no differences in the flux output between treatment conditions. Model 2 regression of the relationship between flux and CFUs for each conditions show overlapping slope and intercept (imd$^{10191}$: y= 0.59 x+ 7.4, imd$^{10191}$+ AMP: G130: y= 0.66 x+ 6.5). This suggests that within this time frame the absence of a plasmid selection does not affect flux output in a significant manner.

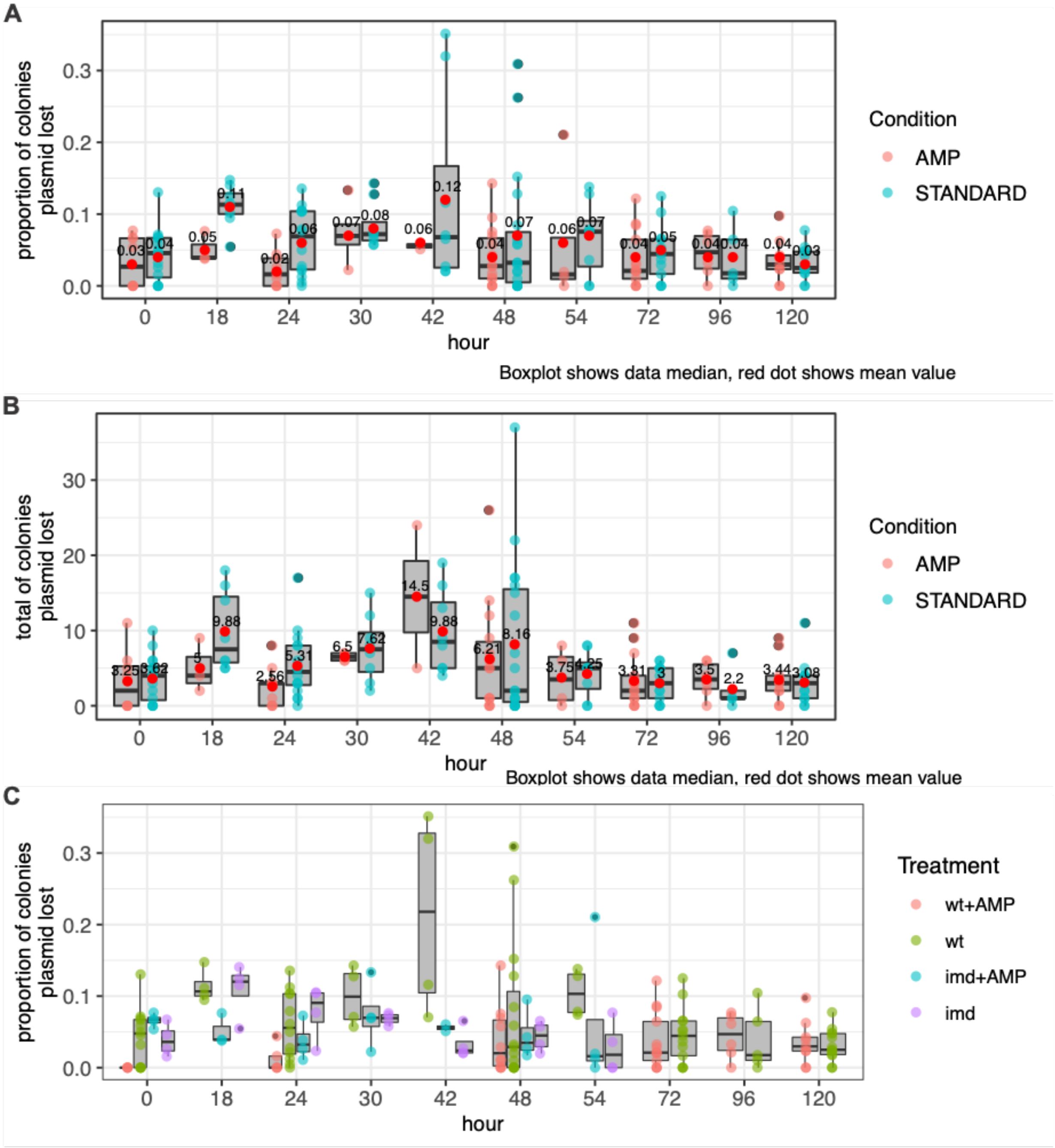


**Figure S6: *ilux-Ecoli* injected into flies show minimal plasmid loss over time.** To ensure *ilux-Ecoli* maintain plasmid expression during time course experiments, we measured the ratio of luminescent vs non-luminescent colonies from injected wild-type (wt) and imd$^{10191}$ flies grown on standard media or media supplemented with the plasmid selective compound ampicillin. Two seperate injection concentrations were used for wt vs imd$^{10191}$ flies. This is because wt flies are able clear the infection and so require a higher starting dose in order for the infection to remain above our detection threshold. Immune deficient flies however, die relatively quickly and so require a low initial dose and more frequent sampling over the course of infection. Thus wt flies were injected with 34 nL of *ilux-Ecoli* at an OD=6 and imd$^{10191}$ flies were injected with 34 nl of ilux-

Ecoli at OD=0.06 . The number of bright and dark colonies from injected flies were measured over the course of 120 hours for wt flies, or over the course of 54 hours for $imd^{10191}$. A) We found over the course of 5 days mean plasmid loss across both genotypes and conditions remained relatively low. B) Total counts of colonies with plasmid loss. C) Proportions of plasmid loss experienced by *ilux-Ecoli* stratified by genotype and treatment. We would anticipate that if plasmid loss was a potentially confounding factor we should observe it most readily in the actively proliferating infection in immune deficient flies. However this does not appear to be the case. There do not appear to be differences in the plasmid loss in infections regardless of the presence of ampicillin, a plasmid selective compound.

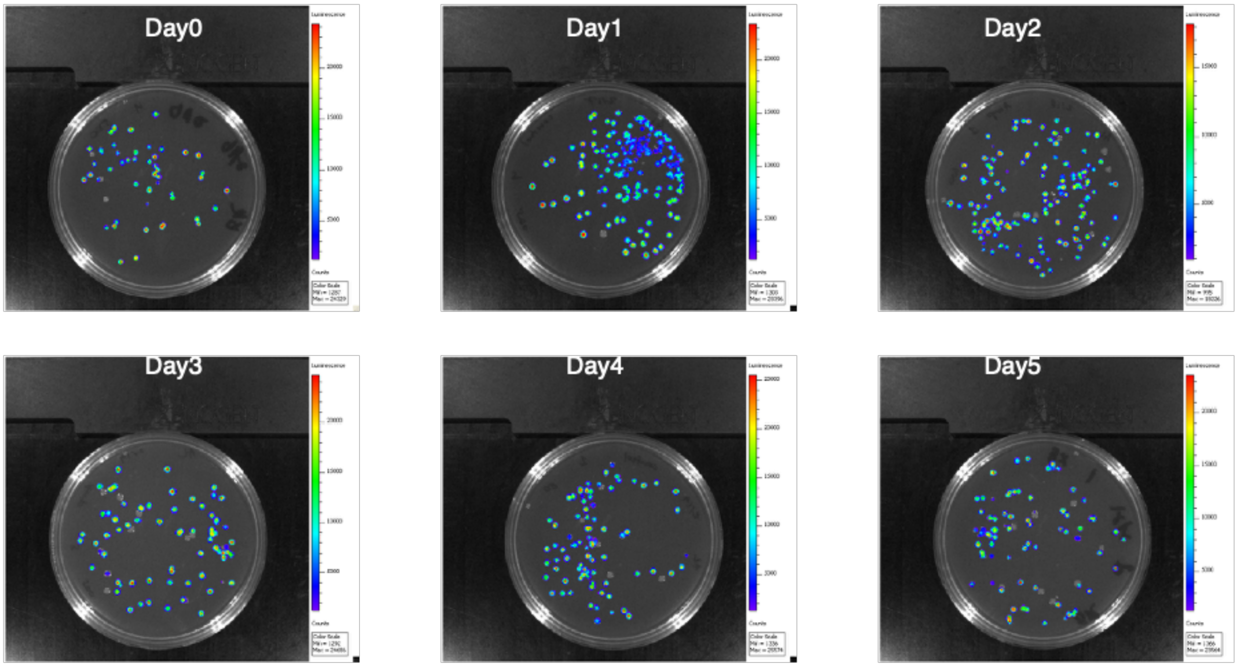


**Figure S7 : Representative images of colonies used to measure plasmid loss.** To assess bright and dark colonies, plates were imaged and colonies assessed for bioluminescence emission. Bright colonies were called as any colonies showing signal above the background level of radiance.

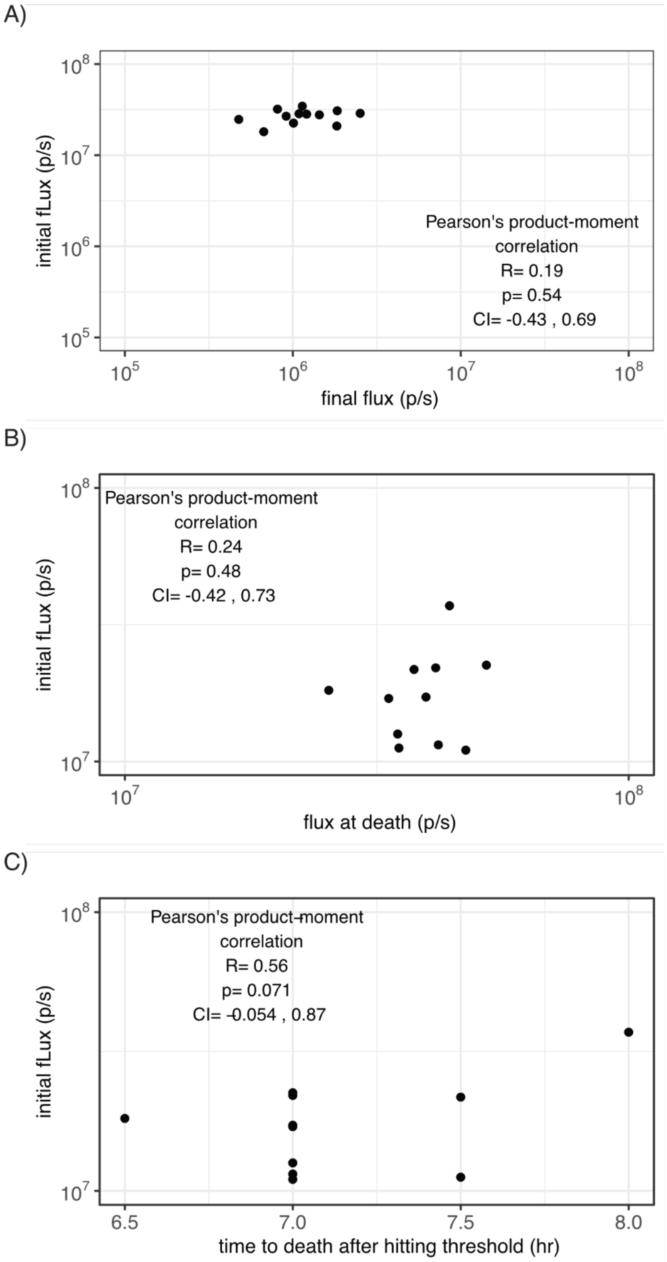


**Figure S8: Initial dose of infection does not result in differences in final load or time to death in Oregon-R or imd$^{10191}$.** A) To determine if differences in the final load of infection may be a result of differences in the initial load, we tested for correlation between flux at the time of injection and flux at the end of the time course in wild-type flies (data shown in Figure 4). We did not find a significant correlation between the two (R= 0.19, p= 0.54, Pearson product-moment correlation). B) Next in the imd$^{10191}$ genotype, we sought to determine if differences in the load of infection upon death may be a result of differences in the initial load. We tested for correlation between flux at the time of injection and flux at the end of the time course and did not find any significant correlation between the two (R = 0.24 , p= 0.48, Pearson product-moment correlation). C) Lastly, we wondered if differences in the initial load of infection may correlate with differences in the time of death in imd$^{10191}$ flies after hitting the mean bacterial load upon death We did not find a significant correlation between these two variables (R= 0.56, p= 0.076, Pearson product-moment correlation).

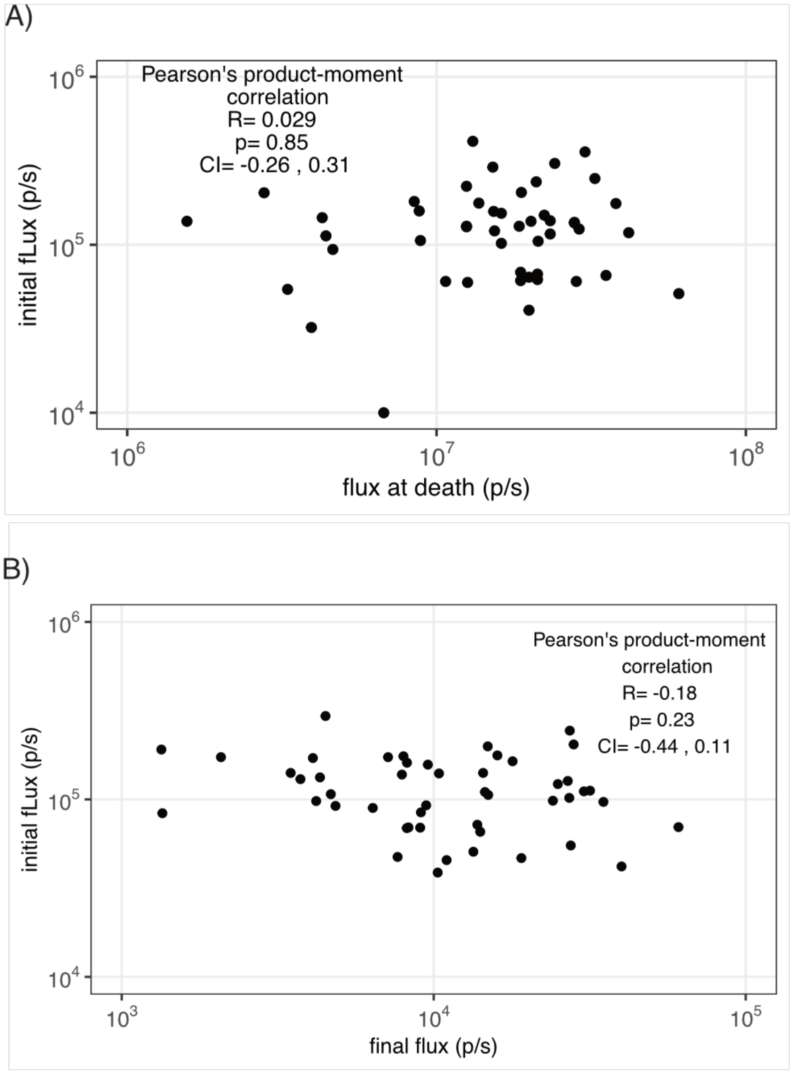


**Figure S9: Initial dose of infection does not result in differences in final load in imd$^{10191}$.** A) Given the higher number of biological replicates (n=48) of this experiment (Figure 5) over the previous (n=12) (Figure 4), we again asked if differences in the initial load of infection for imd$^{10191}$ flies resulted in differences in the radiance upon death. We found no significant correlation between these two variables (R = 0.029, p = 0.85, Pearsons's product-moment correlation). B) To determine if the differences in the initial load of infection could explain differences in the radiance at 48 hours, we tested for correlation between these two variables in wild-type flies. We did not find significant correlation between the two variables (R = -0.18, p = 0.23, Pearsons's product-moment correlation).

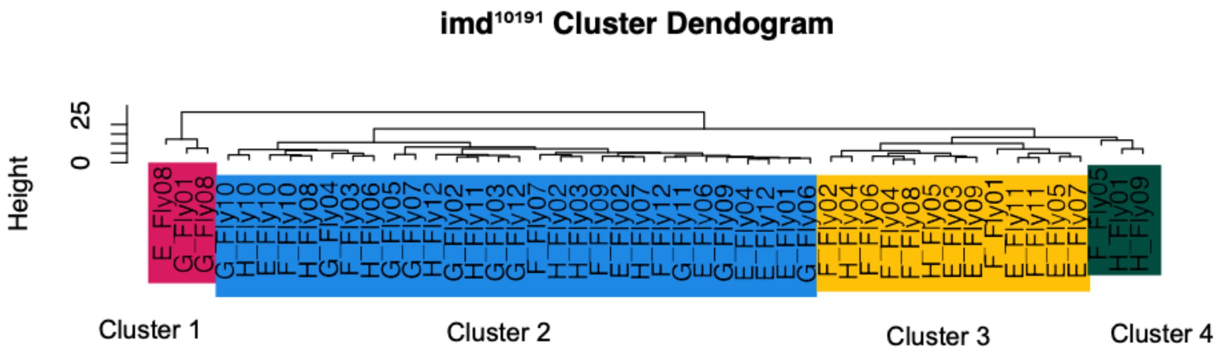


**Figure S10: Unsupervised clustering of imd$^{10191}$ results in 4 distinct clusters.**
In order to group infection profiles we performed hierarchical clustering using Euclidean dissimilarity on log transformed radiance data. Clusters were assigned based on the groups resulting from the first 3 branching points of the dendrogram. For this analysis individual flies were named after their position in the plate with the letter corresponding to a row in the plate and the fly number corresponding with the plate column.

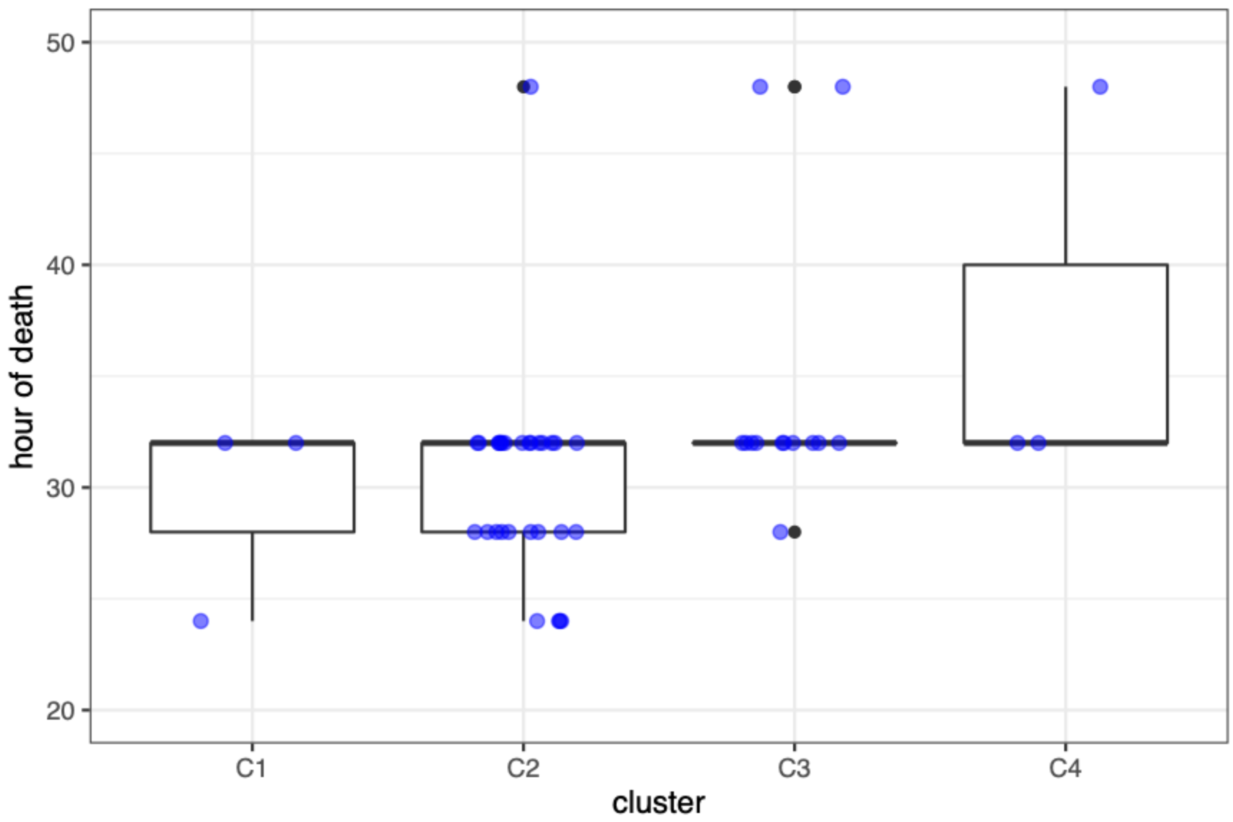


**Figure S11: Time of death does not correlate with assigned cluster.** We asked if there was any cluster that was enriched for a particular time of death. We performed Welch unpaired two sample t-test on every cluster combination and found that no cluster mean appeared to be significantly different from the rest, using a p-value cutoff of 0.05, C1 vs. C2 $p$=0.7796, C1 vs. C3 $p$=0.2054, C1 vs. C4 $p$=0.2739, C2 vs. C3 $p$=0.05494, C2 vs. C4 $p$=0.3123, C3 vs. C4 $p$=0.6186.

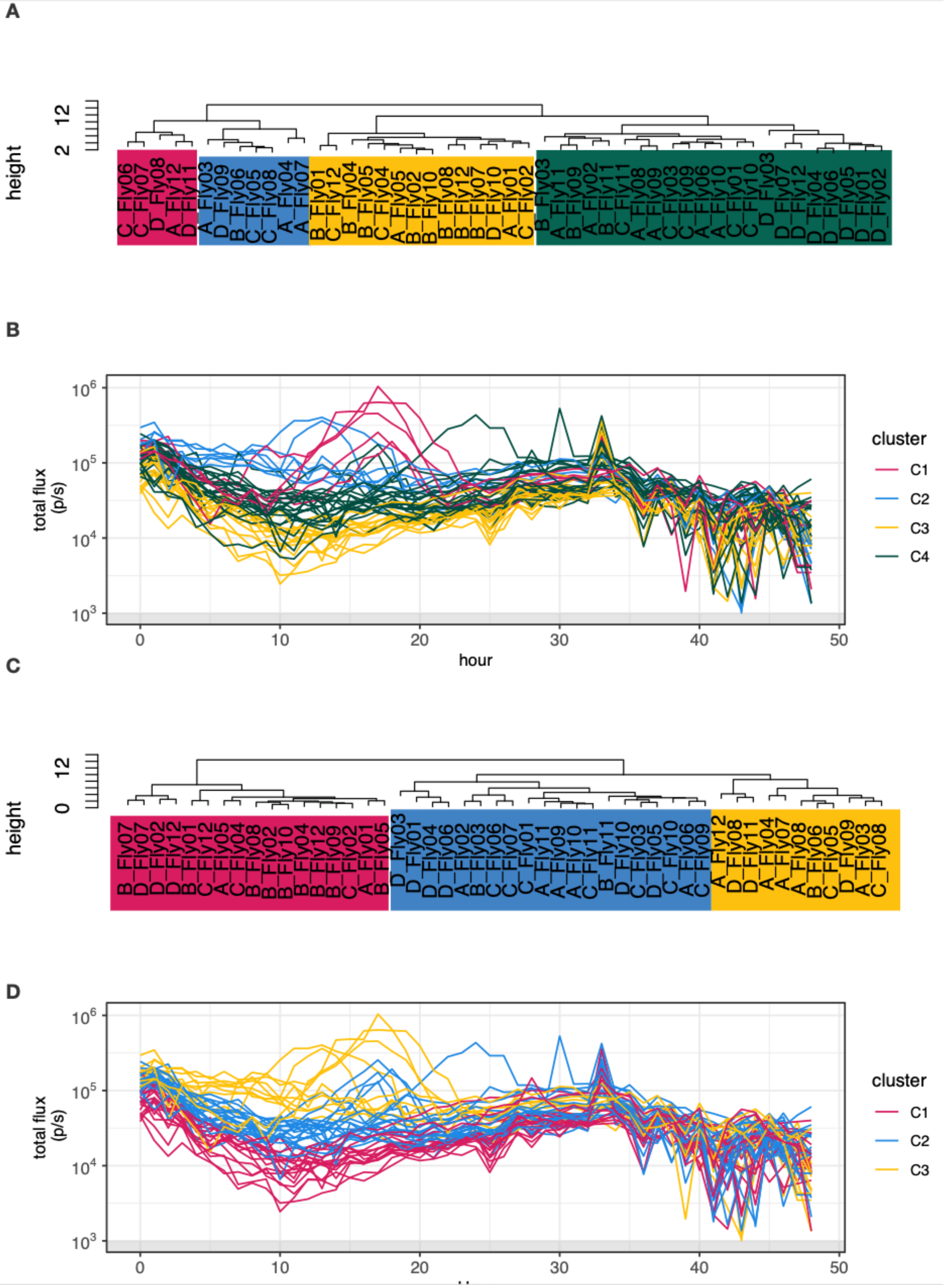
A
height
12
2
B
total flux (p/s)
$10^6$
$10^5$
$10^4$
$10^3$
0
10
20
30
40
50
hour
cluster
C1
C2
C3
C4
C
height
12
0
D
total flux (p/s)
$10^6$
$10^5$
$10^4$
$10^3$
0
10
20
30
40
50
cluster
C1
C2
C3

**Figure S12: Unsupervised clustering of Oregon-R infection profiles does not separate trajectories by infection resurgence.** In order to group infection profiles, we performed hierarchical clustering using Euclidean dissimilarity on log transformed radiance data. We noticed that several flies showed an increase in bacterial load between 10-25 h. A) Clustering using the full data, clusters were assigned based on the first 3 branching points of the dendrogram. For this analysis individual flies were named after their position in the plate with the letter corresponding to a row in the plate and the fly number corresponding with the plate column. B) Coloring of individual infection profiles shows that initial clustering fails to group profiles that show a resurgence in bacterial load. C) We suspected that the general convergence of the data at around hour 35 and the noise after the 40 h mark may be affecting the clustering, so we performed a second round of clustering using only data from before hour 30. D) Coloring of individual infection profiles based on clustering shows that the clustering of the censored data similarly does not group profiles based on our feature of interest.

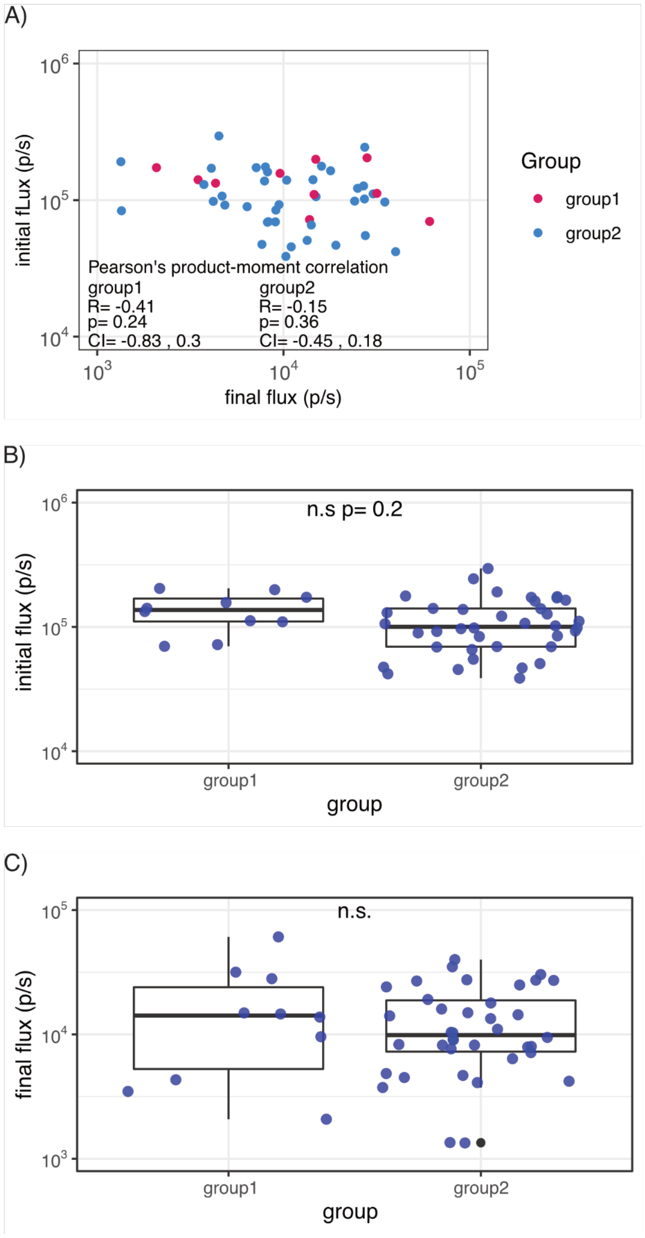


**Figure S13: Initial load of infection does not result in differences in final radiance or resurgence of infection.** A) To determine if the initial load of infection for wild-type Oregon-R flies resulted in differences in the final radiance, we measured the correlation between the initial and final flux for individual flies. We found that the difference in initial dose of infection did not correlate with the final flux in either group (group 1 R=-0.41, p= 0.24, group 2 R=-0.15, p= 0.36, Pearson's product-moment correlation). B) To determine if the resurgence of infection that is characteristic in group 1 could be explained by differences in the initial dose of infection between group 1 and group 2, we compared initial loads between the groups We did not observe a difference in the initial load between these groups (group 1 vs. group 2 initial p=0.2, Welch two sample t-test). C) Lastly we wondered if the resurgent infection observed in group 1 resulted in differences in the measured load of infection at 48 hours. We did not observe a difference in the initial load between these groups (group 1 vs. group 2 p=0.45, Welch two sample t-test).

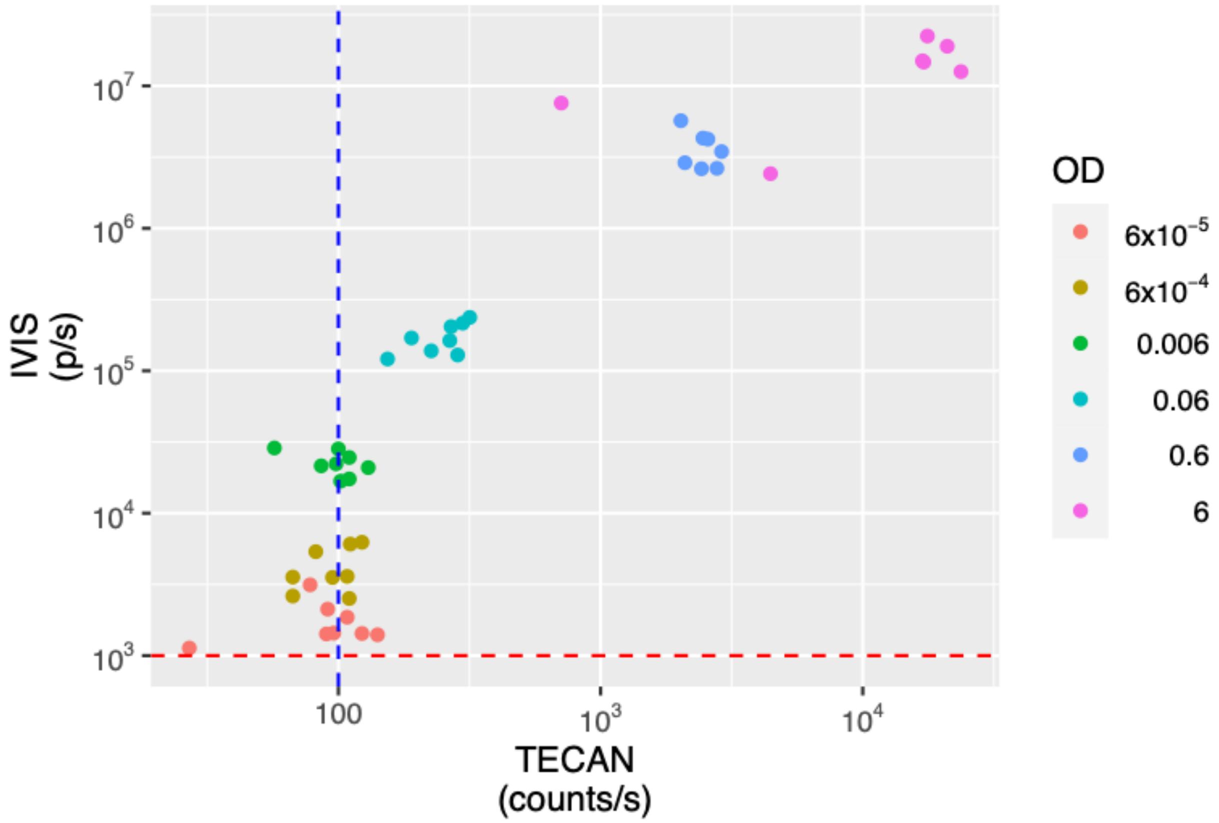


**Figure S14: Autobioluminecent bacteria is detectable by both IVS and plate readers.** To determine how amenable this method is to more cost effective equipment set-ups we compared readings from flies between our set up (IVIS) and a TECAN plate reader. Flies were injected with 6 doses of bacteria ranging from approximately 10,000 CFUs/fly (OD=6) to less than 10 CFUs (OD=6x10$^{-5}$). Thresholds of detection for both equipment are marked as a red dashed line for the IVIS and blue dashed line for the TECAN plate reader. We observe that the IVIS is able to distinguish injection doses above an OD of 6x10$^{-4}$ while the plate reader set up is able to distinguish doses above an OD of 0.006. Thus while the IVIS is more sensitive at the lower range of detection both equipment are able to distinguish doses at above an approximately 100CFUs/fly.
